# Fine-grained descending control of steering in walking *Drosophila*

**DOI:** 10.1101/2023.10.15.562426

**Authors:** Helen H. Yang, Luke E. Brezovec, Laia Serratosa Capdevila, Quinn X. Vanderbeck, Atsuko Adachi, Richard S. Mann, Rachel I. Wilson

## Abstract

Locomotion involves rhythmic limb movement patterns that originate in circuits outside the brain. Purposeful locomotion requires descending commands from the brain, but we do not understand how these commands are structured. Here we investigate this issue, focusing on the control of steering in walking *Drosophila*. First, we describe different limb “gestures” associated with different steering maneuvers. Next, we identify a set of descending neurons whose activity predicts steering. Focusing on two descending cell types downstream from distinct brain networks, we show that they evoke specific limb gestures: one lengthens strides on the outside of a turn, while the other attenuates strides on the inside of a turn. Notably, a single descending neuron can have opposite effects during different locomotor rhythm phases, and we identify networks positioned to implement this phase-specific gating. Together, our results show how purposeful locomotion emerges from brain cells that drive specific, coordinated modulations of low-level patterns.

## Introduction

Vertebrates and arthropods are unique among living creatures in their ability to walk and run. All walking or running organisms confront the same problem — namely, how to keep the body raised above the ground with a stable center of mass, while also propelling the body forward at the intended speed and steering along the intended path. The solution is to use different limbs for propulsion at different moments, in a rhythmic cycle (Figure 1A). Each limb alternates between a power stroke (stance) and a return stroke (swing). In the power phase, the limb is in contact with the ground, where it exerts propulsive force. In the return phase, the limb swings to a new position. Forward speed control emerges from symmetric modulations of this limb movement pattern, whereas steering emerges from asymmetric modulations (Figure 1A).

**Figure 1:**
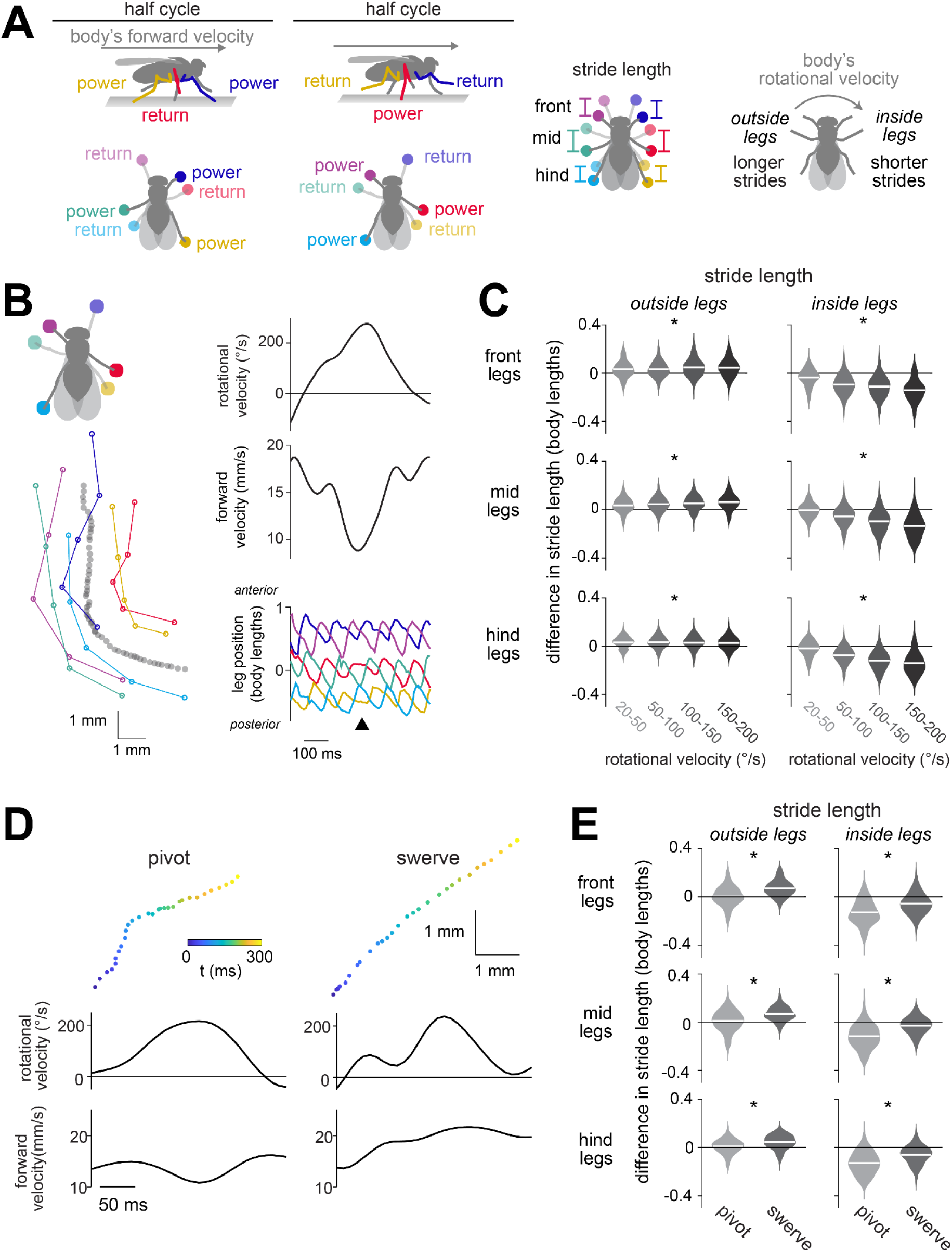
Different steering maneuvers use different leg gestures. (A) During the power stroke, the leg pushes backward. During the return stroke, the leg swings forward to a new position. Schematics viewed from below show each leg at the endpoint of its power and return stroke. Stride length is the difference between the two endpoints in the anterior-posterior axis. (B) *Left*: an example steering bout, showing the path of the fly’s body (gray) and the positions of the leg tips as they touch down at the end of the return stroke of each step. *Right*: the fly’s rotational and forward body velocity during the bout as well as the anterior-posterior position of the leg tips in body-centric coordinates. The arrow highlights the stride length changes at the peak of the turn. (C) When flies rotate, stride length decreases for the inside legs, while increasing for the outside legs. For each turning bout, we measured the bout’s peak rotational velocity, as well as the stride length for the step that occurred at that peak. Stride lengths are expressed as the change from the mean when the fly was not turning. Stride length varies significantly with rotational velocity for both outside and inside legs (two-way ANOVA, rotational velocity and legs as factors, rotational velocity: p=4.87×10^−77^; interaction between rotational velocity and leg identity: p=0); post-hoc Tukey-Kramer tests show significant changes for every leg at all but the lowest rotational velocity (see Table S1)), n = 138 (55), 454 (68), 434 (68), and 151 (50) bouts (flies) for 20-50, 50-100, 100-150, and 150-200 °/s, respectively. * marks legs with significant changes during turning (D) *Top:* paths for example steering bouts, corresponding to a pivot and a swerve (downsampled to 100 positions/s). *Bottom:* rotational and forward velocity over time for these examples. (E) Pivots and swerves produce different changes in stride length (two-way ANOVA with legs and pivot/swerve as factors: pivot/swerve p=6.11×10^−31^; leg identity p=4.96×10^−66^; interaction: p=0.738, n = 389 (64) and 155 (53) bouts (flies), respectively). Post-hoc Tukey-Kramer tests show a significant difference between pivots and swerves for every leg (see Table S1).

These modulations require cooperation among several regions of the central nervous system. Specifically, the rhythmic limb movement pattern is intrinsic to the vertebrate spinal cord^1,2^ or the arthropod ventral nerve cord^3–7^. Meanwhile, descending signals from the brain to the cord are required to start or stop locomotion, modulate speed, or change direction in both vertebrates and arthropods^8–12^. Yet the brain’s role in locomotor control is not necessarily limited to general high-level commands: blocking descending signals from the brain can also alter the details of limb coordination during walking in both vertebrates^13–15^ and arthropods^7,16^, and stimulation of specific descending tracts can produce excitation or inhibition of muscle groups in multiple limbs^17–21^. Currently, we do not understand what features of locomotion are actually under the brain’s control.

In considering this problem, it is useful to consider the concept of elemental components. In a puppet, the elemental components of control are the strings, and there are fewer strings than joints because the puppeteer is simply concerned with evoking naturalistic gestures, not arbitrarily controlling every joint; similarly, the brain may have a limited number of control “strings”^22^. Circuits in the spinal cord or ventral nerve cord can control sets of co-activated muscles, and so these muscle synergies are often treated as the elemental components of control. However, from the brain’s perspective, the elements of motor control might be more high-level than these muscle synergies. Once we can map the brain’s strings onto specific descending cell types, we should be in a better position to understand the logic of control.

In this regard, *Drosophila* offers unique advantages: it is the only limbed animal with a near-complete connectome^23–27^, and many *Drosophila* descending neurons (DNs) involved in walking control are uniquely identifiable as individual right-left cell pairs^28–34^. Moreover, new genetic tools^28,35^ allow us to target single DNs for recording and perturbation during locomotion.

In this study, we use walking *Drosophila* to focus specifically on the logic of steering control, as steering is a universal and complex problem. Steering must arise from right-left asymmetries in limb movements in walking *Drosophila*^36,37^ and other hexapods^38–46^, just as in tetrapods and bipeds^2,47^. In all these species, steering is accomplished using a variety of limb movement patterns, all requiring limb specialization and coordination. Currently, we know relatively little about the descending control of steering in any species, aside from the fact that descending signals are asymmetric during a turning bout^20,30,31,48–55^. A recent study found that multiple DN axons in the neck of a walking fly were correlated with steering direction but not with limb movement variations during steering^56^; however, it is curious that there should be multiple DN axons driving steering if all these cells are merely carrying redundant copies of the same general command to turn right or left.

Our study has three parts. First, we identify different limb movement gestures associated with different types of steering maneuvers. Second, we identify a set of DNs whose activity is correlated with steering; this provides a potential neural substrate for the elemental control of limb gestures. Third — in the main focus of the study — we show that different steering DNs evoke distinct limb gestures and that they are recruited by essentially non-overlapping pathways in the brain. Moreover, we demonstrate that a single DN can have opposite effects on leg movement during different phases of the locomotor cycle, and we identify cells in the ventral nerve cord that are anatomically positioned to implement this phase-specific gating. Together, our results show how the brain can exert detailed but also efficient control of steering via selective recruitment of DNs dedicated to specific limb gestures. Our findings help re-cast the abstract problem of adaptive locomotor control as a more concrete problem of generating specific sequences of activity in populations of neurons in the brain.

## Results

### Different steering maneuvers use different leg gestures

To understand the descending control of steering, we first need to understand what variations in steering actually look like. It is known that *Drosophila* can perform several types of steering maneuvers, which are associated with distinct changes in the body’s rotational and forward velocity^57,58^. Steering can also involve several different leg “gestures”^36,37^, defined here as coordinated changes in the movement of multiple limbs (Figure 1A). We wondered whether different types of steering maneuvers use systematically different leg gestures; if so, then these multi-leg gestures might be the relevant elemental components of descending control.

When we analyzed the leg movements of freely walking flies^59^, aided by high-resolution, automated body-part tracking^60^ (Figures 1B and S1A), we found that three asymmetrical, multi-leg gestures were clearly associated with steering: increasing the stride length of the three legs on the outside of the turn (Figure 1C), decreasing the stride length of the three legs on the inside of the turn (Figure 1C), and shifting the step direction of the front legs so as to pull the body into the turn (Figure S1B). It should be emphasized that stride length modulations during steering are very small, on the order of 100 μm, or 5% of the body length; nonetheless, they can produce fast body rotations because they are repeated at a rate of about 10 Hz and are deployed in combination. This overall description of leg movements during steering is largely consistent with previous work^36,37^.

Supporting our hypothesis, we found that different types of steering maneuvers were associated with systematically different leg gesture usage. Specifically, when we divided steering maneuvers into two types, pivots and swerves, we found a systematic difference in these leg gestures. We define a “pivot” as an epoch where the body rotates while slowing or reversing forward movement. Conversely, we define a “‘swerve” as an epoch where the body rotates while increasing forward movement. Whereas a pivot tends to produce tight kinks in a walking path, a swerve produces a smooth curve (Figure 1D). We found that the first gesture (increasing stride length on the outside of a turn) was more prominent in swerves versus pivots, whereas the second gesture (decreasing stride length on the inside of a turn) was more prominent in pivots versus swerves (Figure 1E). These differences in stride length resulted from changes in the power stroke as well as changes in the return stroke (Figure S1C). We also found that the third gesture (shifting step direction) was more prominent in pivots versus swerves (Figure S1C).

In short, different steering maneuvers arise from significantly different combinations of leg gestures. This finding suggests that leg gestures are elemental components of neural control — that is, they are among the fundamental building blocks used to generate a particular path through the environment. Given this, we wondered whether the brain can independently control the recruitment of different leg gestures. Therefore, we next set out to expand the list of identifiable DNs associated with steering control, with the ultimate goal of determining whether any of these cell types are responsible for generating specific leg gestures.

### Many descending neuron types correlate with the body’s rotational velocity

In order to expand the list of identifiable steering-related DNs, we focused on DNs that are genetically accessible, meaning that there exists a specific transgenic driver line that can be used to target that DN for perturbation or recording^28^. We specifically examined those DN types that are anatomically positioned to influence at least two legs, meaning that they project to at least two homolateral (“same-side”) leg neuromeres in the ventral nerve cord. This pointed us toward 16 DN types. Here we use “DN type” to refer to a morphological category of DNs^28^. Most DN types innervating the leg neuromeres consist of a single right-left cell pair^28^, and this was true of all the DN types we examined in this study.

In each DN type, we co-expressed the genetically encoded calcium indicator jGCaMP7s and the red fluorophore CyRFP1 (for motion correction), and we monitored neural activity as the fly walked on a spherical treadmill. We found that all but three of these DN types had increased activity during walking bouts, as compared to non-walking epochs (Figure S2). One of these (DNg34) also showed a graded relationship with forward velocity (Figure S2).

Most relevant to this study, we discovered that five of the DN types in our screen showed right-left activity differences that were consistently correlated with the body’s rotational velocity during steering. Four of these five cell types were correlated with ipsiversive steering—i.e. the cell with a soma in the right hemisphere was more active during right turns and *vice versa* (DNa01, DNa02, DNb05, and DNg13; Figure 2A). Meanwhile, the fifth cell type was correlated with contraversive steering (DNb06; Figure 2A). Previously, only DNa01 and DNa02 were known to be consistently correlated with rotational velocity^30,31^, so our findings substantially expand the list of identified steering-related DNs. For all five DN types, we found that right-left differences in DN activity were linearly correlated with the body’s rotational velocity (Figure 2B).

**Figure 2:**
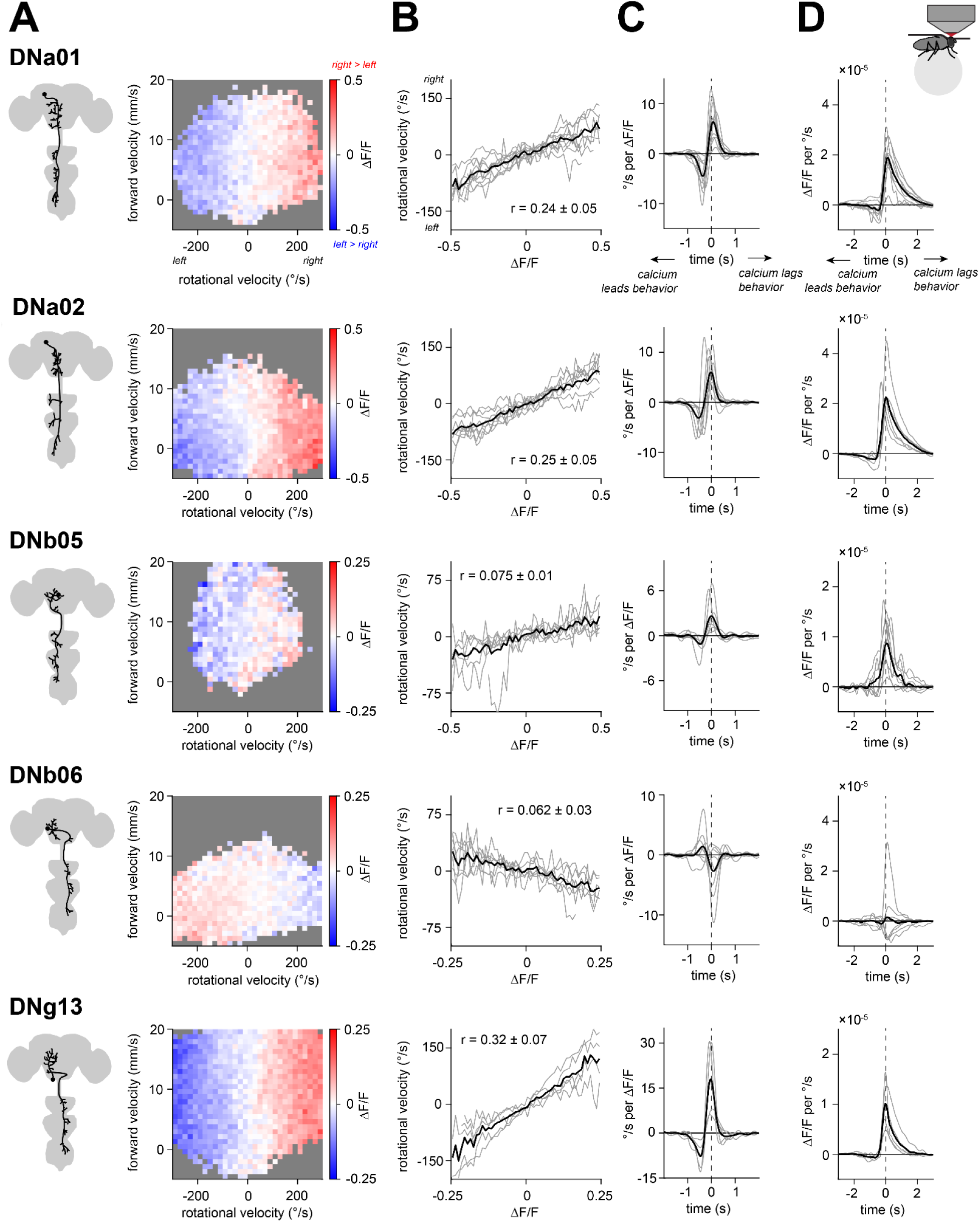
Many descending neuron types correlate with the body’s rotational velocity. DNa01, DNa02, DNb05, DNb06, and DNg13 are DN types represented by a single right-left cell pair; here we use calcium imaging to show that all 5 are related to rotational velocity. Figure S2 shows data for other DN types. (A) *Left:* schematic of the left cell for each DN type. *Right:* in example flies, right-left difference in ΔF/F, shown as a heatmap over rotational and forward velocity bins. We allocated each time point to a bin (excluding times when the fly was not walking) and took the mean right-left difference in ΔF/F within each bin. Gray bins had <20 time points. (B) Rotational velocity is related to the right-left difference in ΔF/F. We allocated each time point to a velocity bin (excluding times when the fly was not walking) and took the mean right-left difference in ΔF/F within each bin. Individual flies are in gray, with the mean across flies in black. The mean ± SEM across flies of the correlation between rotational velocity and ΔF/F is shown. For (B-D), n (flies) is 10 (DNa01), 8 (DNa02), 7 (DNb05), 7 (DNb06), 5 (DNg13). (C) Filters describing the mean rotational velocity around an impulse change in the right-left difference in ΔF/F. Here and in (D) we did not exclude times when the fly was not walking. (D) Filters describing the mean right-left difference in ΔF/F around an impulse change in rotational velocity.

If these DNs influence steering, we would expect that their activity precedes changes in rotational velocity. To investigate this question, we computed the linear filters relating rotational velocity to neural activity, where “neural activity” here specifically means the right-left difference in ΔF/F in a given DN type. We can compute a linear filter in two ways. First, we can compute a neuron→behavior filter that describes the average relationship between a transient step of neural activity (a unit impulse) and the associated change in rotational velocity (Figure 2C). Conversely, we can compute a behavior→neuron filter that describes the average relationship between a unit impulse of behavior and the associated change in neural activity (Figure 2D). These filters are computed in a manner that accounts for the limited speed of calcium signals as well as the characteristic timescales of behavior. For both types of filters, we found that changes in ΔF/F preceded changes in behavior for all of the turning-correlated DNs (Figures 2C and 2D). These results are compatible with the idea that these DNs are causal for steering.

To recap, our data point to five specific DN types whose activity precedes and predicts steering. At the level of calcium imaging, these five cell types appear to carry redundant signals. However, it seems unlikely that all these DNs are merely carrying redundant copies of the same rotational velocity command. Rather, we wondered whether different steering DNs might have specialized roles in fine-grained control of leg movements.

### Ipsiversive steering-correlated descending neurons DNa02 and DNg13 have distinct inputs and outputs

To test this hypothesis, we decided to focus on DNg13 and DNa02, which had the strongest correlation with rotational velocity (Figure 2B). From the perspective of their soma and dendritic arbor, both DN types predicted ipsiversive steering. However, DNg13 axons cross the midline as they descend into the ventral cord, whereas DNa02 axons do not. Furthermore, when we used the whole brain connectome^23,27^ to identify the direct inputs to DNa02 and DNg13, we found that these two cells receive input from largely non-overlapping individual presynaptic neurons (Figures 3A, S3A, and S3B). This finding implies that DNa02 and DNg13 can be recruited independently.

**Figure 3:**
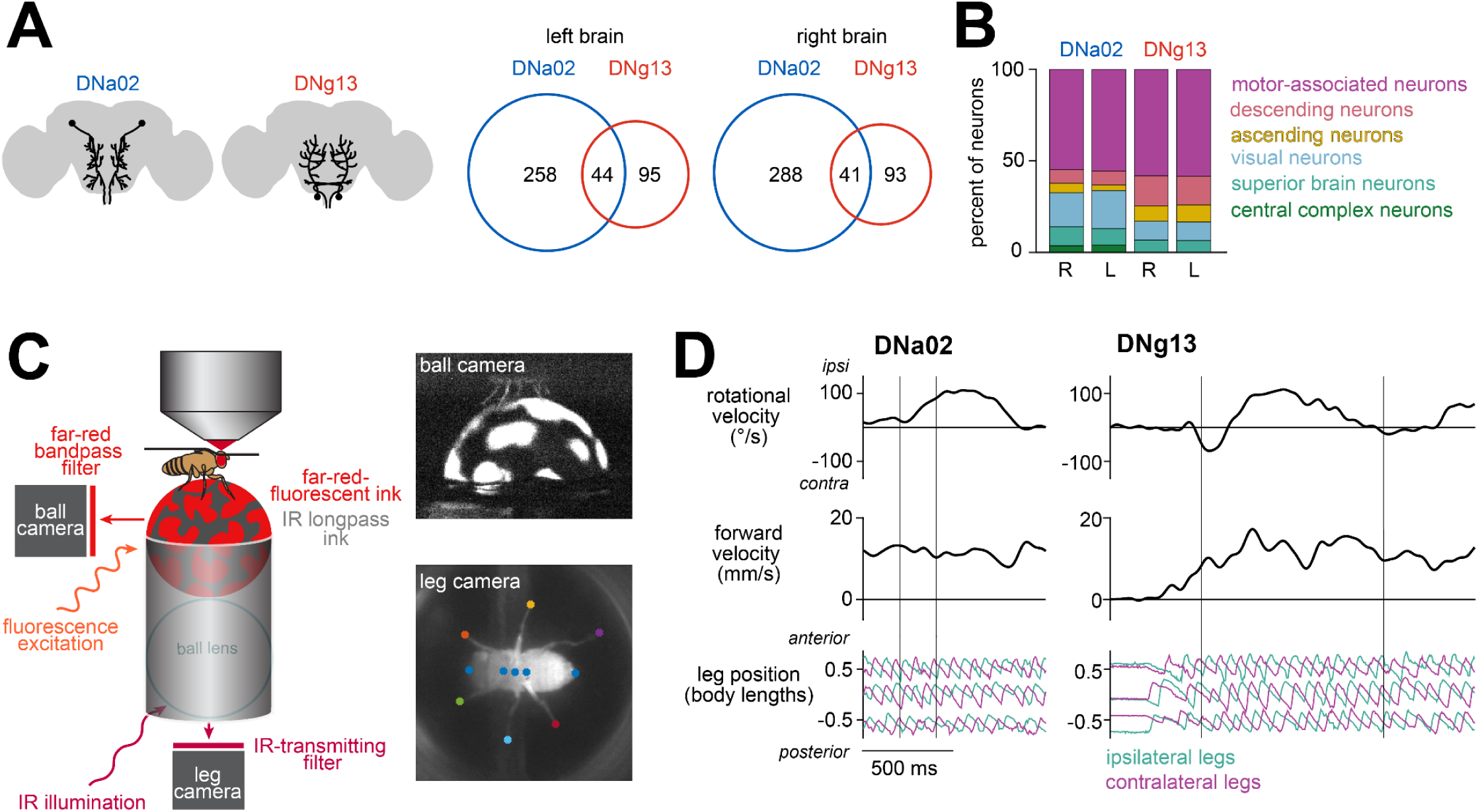
DNa02 and DNg13 have distinct inputs in the brain and distinct effects on leg movement. (A) *Left:* Schematic of the left and right DNa02 and DNg13 cells in the brain. *Right*: Analysis of the full brain connectome shows that DNa02 and DNg13 neurons in the same hemisphere share relatively few presynaptic cells (numbers are cell counts). (B) Stacked bar charts summarize the cells presynaptic to each DN in the brain, divided into 6 categories, and expressed as a percentage of all presynaptic neurons (again, in the brain). (C) The fly walks on a spherical treadmill coated with IR long-pass ink and painted with spots of far-red fluorescent ink. The camera used for tracking the sphere (and thus the fly’s fictive body velocity) captures only far-red light, making the ball appear black with white spots (*top*). The camera used for tracking the legs captures only IR light; because the sphere is transparent to IR light, it is not visible in this view (*bottom*). The key points, as tracked by APT, are marked. (D) Example trials for unilateral optogenetic stimulation of DNa02 with CsChrimson (*left*) and unilateral depolarization of DNg13 by current injection (*right*). Thin lines mark the onset and offset of stimulation.

An interesting example of this is the pattern of projections from the central complex, the locus of spatial memory and navigation in the insect brain. DNa02 is directly targeted by central complex output neurons responsible for feedback control of heading^31,61–63^. By contrast, DNg13 receives no direct output from the central complex.

Aside from the central complex, we did not find clear evidence of any regions being upstream from just one of these cells. For example, we found that both DNs are downstream of visual regions (the optic lobe and optic glomeruli). Moreover, both DNs are postsynaptic in motor-associated brain regions. Both DNs also receive substantial input from other DNs; this is not surprising, as many DNs communicate via specific networks of dendro-dendritic connections in the brain^64^. Finally, both DNs receive input from ascending neurons. Thus, these DNs participate in similar categories of brain networks; they are simply targeted by different sets of presynaptic cells.

### Different descending neurons drive distinctive leg gestures

To determine how each DN influences steering, we set out to perturb neural activity unilaterally. In each experiment, we simultaneously monitored the positions of all six legs, as well as the body’s fictive velocity. Specifically, we built a spherical treadmill system that could appear either transparent or opaque, depending on the wavelength of illumination (Figure 3C). This system allowed us to image the legs from below via light transmitted straight through the spherical treadmill while also imaging the surface of the sphere itself via a separate optical pathway, allowing us to infer the fly’s fictive body velocity as it walked on the sphere. We monitored these behavioral parameters as we perturbed the DNs (Figure 3D).

To unilaterally stimulate DNa02, we expressed the light-gated cation channel CsChrimson in individual DNs using a genetic mosaic strategy^35^. We compared the behavioral responses of experimental flies (having expression in one copy of DNa02) with control flies (having expression in neither copy of DNa02); note that these flies are genetically identical and have the same average pattern of expression in “off target” cells. In experimental flies, we found that a brief pulse of light (200 ms) triggered a turn in the direction of the stimulated cell, with no change in forward velocity as compared with controls (Figure 4A). Bilateral activation of DNa02 (in flies where the mosaic strategy resulted in expression in both copies) instead triggered a decrease in forward velocity without a change in the heading direction (Figure S4A).

**Figure 4:**
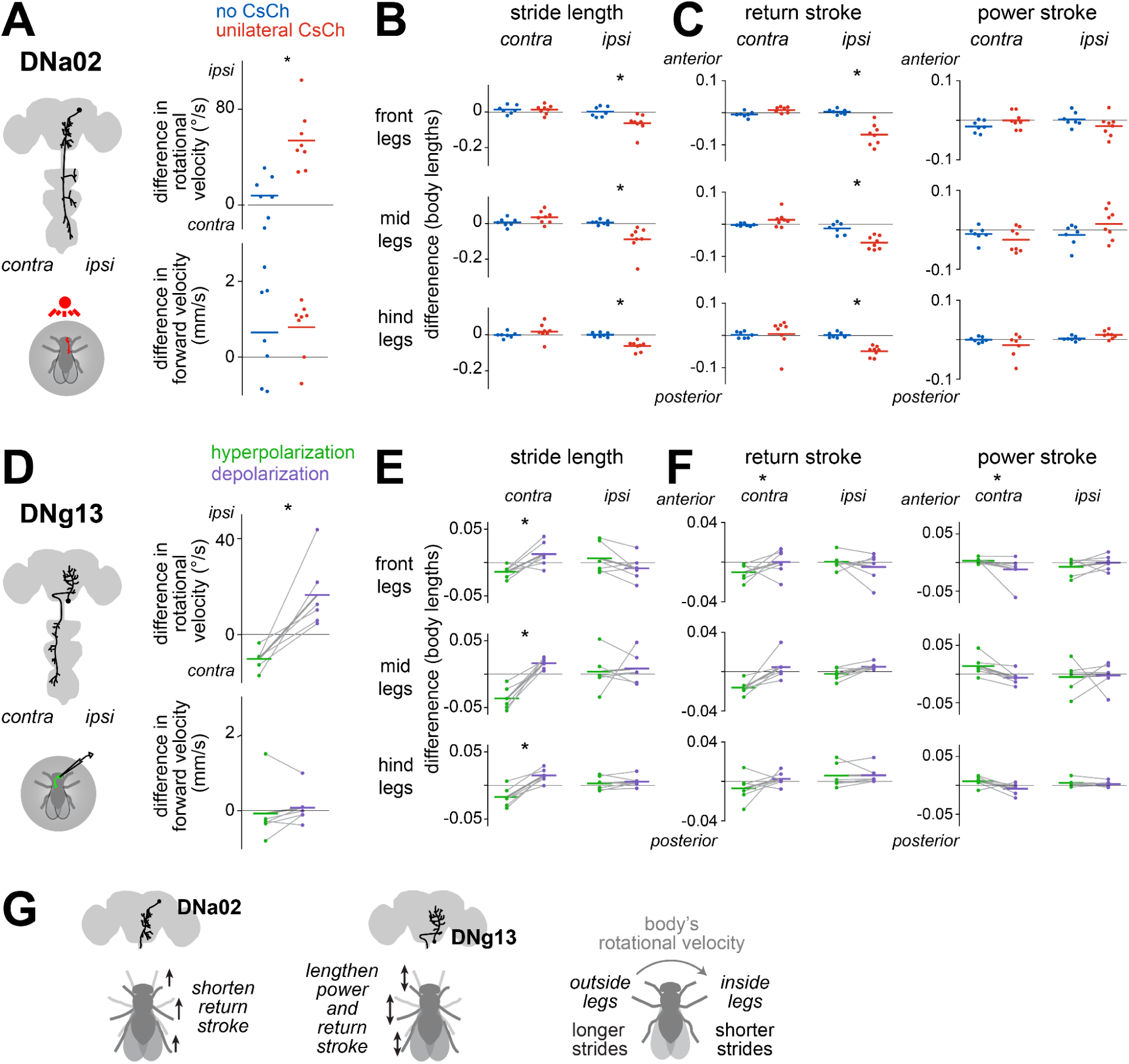
Different descending neurons drive distinctive leg gestures. (A) Difference in rotational and forward velocity during unilateral optogenetic stimulation of DNa02 with CsChrimson (CsCh). Dots are individual flies, and line is the mean across flies. Ipsiversive (positive) and contraversive (negative) are relative to the side of the soma of the CsCh+ cell; for no CsCh controls, ipsiversive and contraversive were randomly assigned for each fly. The effect of optogenetic stimulation is significant for rotational velocity (p=0.00141) but not forward velocity (p=0.802; two-sample t-tests, n=8 flies (CsCh+) and n=7 flies (no CsCh)). * marks changes upon stimulation. (B) Unilateral optogenetic stimulation of DNa02 produces leg-specific changes in stride length (two-way ANOVA with CsCh expression and legs as factors, ±CsCh: p=2.85×10^−4^; interaction between leg identity and ±CsCh: p=8.78×10^−6^, with significant effects for ipsilateral but not contralateral legs in post-hoc tests, as shown in Table S1). (C) Unilateral optogenetic stimulation of DNa02 shortens the return stroke but not the power stroke (return stroke: p-values for two-way ANOVA are 4.23×10^−6^ (±CsCh), 2.46×10^−8^ (interaction between leg identity and ±CsCh), with significant effects in ipsilateral but not contralateral legs in post-hoc tests (Table S1). power stroke: two-way ANOVA p=0.807 (±CsCh) and p=0.0354 (interaction between leg identity and ±CsCh)). (D) The effects of DNg13 current injection are significant for rotational velocity (hyperpolarization p=0.00257, depolarization p=0.0404) but not forward velocity (hyperpolarization p=0.828, depolarization p=0.695; two-sample t-tests comparing against no stimulation periods, n=6 flies). Lines connect data points from the same fly. (E) DNg13 current injection produces leg-specific changes in stride length (3-way ANOVA with depolarization/hyperpolarization, leg identity, and fly identity as factors: depolarization/hyperpolarization p=1.49×10^−4^, interaction between leg identity and depol/hyperpol p=0.000501), with significant effects in contralateral but not ipsilateral legs in post-hoc tests (Table S1). (F) DNg13 current injection affects stride length during both phases of the stride cycle in the contralateral legs (return stroke: 3-way ANOVA depolarization/hyperpolarization, leg identity, and fly identity as factors, without interaction, using contralateral legs only, effect of depolarization/hyperpolarization is significant at p=7.84×10^−4^; power stroke: same with p=5.28×10^−4^). (G) Schematic summarizing the effects of stimulating each DN type.

When we examined leg movements during these steering events, we found that stimulating a single copy of DNa02 decreased stride length in all three legs ipsilateral to the stimulated cell, with no effect on the contralateral legs (Figure 4B), consistent with the anatomy of this DN. The effect of DNa02 on stride length was specific to the leg’s return stroke, which was significantly attenuated, while the power stroke was unaffected (Figure 4C). Bilateral stimulation produced the same leg movement changes, except bilaterally instead of unilaterally (Figures S4B and S4C).

Next, we set out to stimulate DNg13 cells. We found we could not use our genetic mosaic strategy to produce specific unilateral stimulation of this cell type (see Methods), so we instead targeted DNg13 via whole-cell patch-clamp recordings. We injected brief pulses of depolarizing or hyperpolarizing current (1 s in duration), thereby increasing or decreasing the cell’s firing rate. These perturbations produced no change in forward velocity relative to controls (Figure 4D). However, we found that depolarizing current triggered a change in heading in the direction of the stimulated cell, whereas hyperpolarizing current triggered a change in heading in the opposite direction (Figure 4D). This result provides further support for the idea that it is the right-left difference in DN firing rates that produces body rotation.

When we examined leg movements, we found that DNg13 perturbations specifically affected the legs contralateral to the stimulated cell, again consistent with the cell’s anatomy. In particular, we found that contralateral strides were significantly lengthened during depolarization as compared to hyperpolarization; this was not true of ipsilateral strides (Figure 4E). This effect on stride length was a consequence of changes in the power stroke as well as the return stroke: leg displacement was significantly increased during both phases of the step cycle (Figure 4F).

Together, our results imply that each of these DNs generates specific leg gestures associated with steering (Figure 4G). DNa02 shortens steps on the inside of a turn, while DNg13 lengthens steps on the outside of a turn. The lateralized specializations of these DNs are intuitively related to their anatomy: both cells drive ipsiversive rotations, but DNa02 projects ipsilaterally, whereas DNg13 projects contralaterally.

Finally, it is worth remembering that a unilateral projection to the ventral nerve cord can influence both sides of the body, via local connections in the cord that coordinate leg movements across the body midline. Indeed, we observed that unilateral stimulation of either DNa02 or DNg13 shifted the step direction of both front legs so as to pull the body into the turn (Figure S4D). In short, each DN produces a coordinated set of leg kinematic changes, comprising homolateral stride length changes and bilateral, leg-specific step direction changes.

### Descending neuron activity is correlated with specific leg gestures

If DNa02 and DNg13 each control specific leg gestures, then we would expect that their activity correlates with and precedes those leg gestures. To test this idea, we performed whole-cell patch-clamp recordings from these DNs in walking flies (Figure 5A). This approach provides better temporal resolution than calcium imaging, and so it is a better match to the fast timescale of leg movements. During each recording, we monitored the positions of all six legs, as well as the body’s fictive velocity on the spherical treadmill (Figure 3C). We observed that the firing rate in both DNs was higher when the fly was walking (Figure S5A) and that it was positively correlated with the body’s rotational velocity in the ipsiversive direction (Figure 5B).

**Figure 5:**
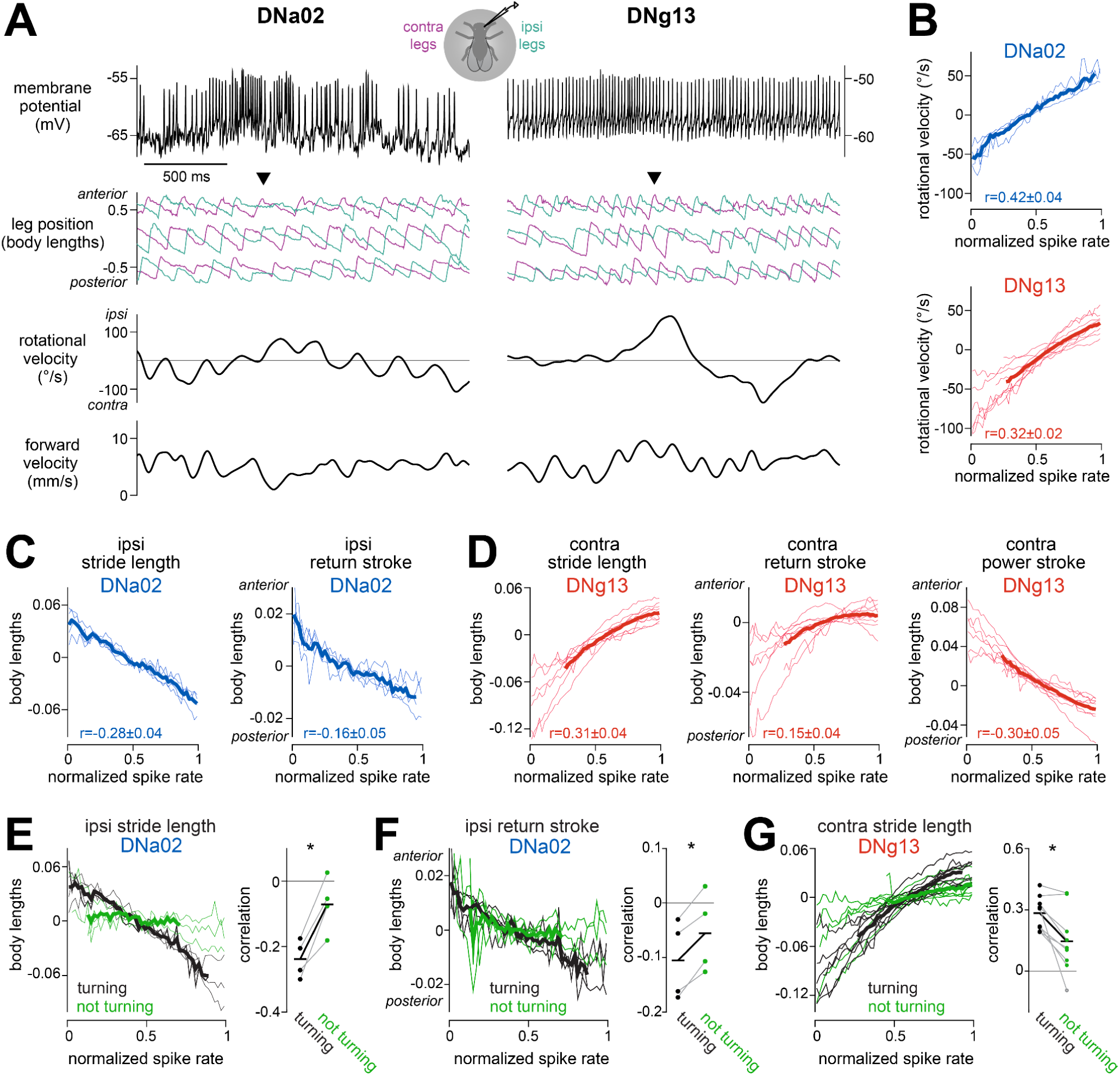
Descending neuron activity is correlated with specific leg gestures. (A) Example recordings. DNa02 spiking increases just before a decrease in ipsilateral stride length, accompanied by shortening of the ipsilateral return stroke and ipsiversive turning (arrow). DNg13 spiking increases just before an increase in contralateral stride length and ipsiversive turning (arrow). (B) Rotational velocity is related to DNa02 and DNg13 spike rate (normalized per cell; see Method details). Positive rotational velocity is ipsiversive turning. Thin lines are mean values for individual flies; thick lines are averages across flies (DNa02: n=4 cells in 4 flies; DNg13: n=9 cells in 9 flies). The mean ± SEM across flies of the correlation between rotational velocity and spike rate is also shown. (C) Same but for leg movement parameters that change with DNa02 optogenetic stimulation (ipsilateral stride length and ipsilateral return stroke position). (D) Same but for leg movement parameters that change with DNg13 current injection (contralateral stride length, contralateral return stroke position, and contralateral power stroke position). (E) The correlation between ipsilateral stride length and normalized DNa02 spike rate only emerges when the fly is turning (defined as rotational speed >25°/s). Summary plot shows the Pearson correlation coefficient between ipsilateral stride length and normalized DNa02 spike rate in both conditions; here pairs of connected dots are flies, and horizontal lines are the means across flies (paired t-test on difference in correlation coefficients, turning versus not turning, p=0.0109). (F) Same as (E) but showing that the correlation between ipsilateral return stroke position and normalized DNa02 firing rate only emerges when the fly is turning (paired t-test on difference in correlation coefficients, turning versus not turning, p=0.00836). (G) Same but analyzing the correlation between contralateral stride length and normalized DNg13 spike rate (paired t-test on difference in correlation coefficients, turning versus not-turning, p=0.00767).

When we analyzed leg movements, we found that DNa02 spike rate was inversely related to ipsilateral stride length, including the ipsilateral return stroke (Figure 5C). This fits with our finding that direct DNa02 stimulation shortens ipsilateral stride length (Figure 4B) through shortening the ipsilateral return stroke (Figure 4C). Together, these results support the conclusion that DNa02 is normally recruited to produce these modulations in leg movement.

Meanwhile, DNg13 spike rate was positively related to contralateral stride length, including both the power stroke and the return stroke of the contralateral legs (Figure 5D). This fits with our finding that DNg13 stimulation increases contralateral stride length, with effects on both the power stroke and the return stroke. Together, these results support the conclusion that DNg13 is normally recruited to produce these modulations.

At the same time, we found that DNa02 and DNg13 were correlated with additional leg kinematic changes that were not observed after direct stimulation of these DNs (Figures S5B and S5C). This result is not surprising because many leg kinematic features are themselves correlated (Figure S5D). Because of this, correlation is not strong evidence of causation in this system. However, a failure to find a correlation would be evidence against causation, which is why it is important to check that DN spiking actually correlates with the expected leg kinematic changes during walking.

Thus far, we have emphasized the fact that asymmetric changes in stride length can rotate the body; however, symmetric changes in stride length can also be used to change the body’s forward velocity^37,65,66^, and symmetric activation of DNa02 did indeed change the stride length and the forward velocity (Figures S4A-C). We therefore wondered whether these DNs might also function as speed-control DNs in addition to functioning as steering DNs. If this were true, then we would expect to find that these DNs are correlated with leg movements whenever the fly is walking, regardless of whether it is turning. In fact, this was not the case: during straight walking, both DNs were much more weakly predictive of their respective step parameters, as compared to epochs where the body was rotating (Figures 5E-G). This result implies that these DNs do not play a major role in the control of forward velocity during straight walking.

### Different descending neurons are recruited at different moments within a steering maneuver

Next, we compared firing rate dynamics in these DNs with behavioral dynamics. On average, firing rate changes in these DNs preceded changes in rotational velocity and stride length by about 150 ms (Figure 6A). This result is consistent with the hypothesis that these DNs actually influence these behaviors. The latency of the behavioral response may partly be due to neural and muscular delays and partly to the inertia of the spherical treadmill itself.

**Figure 6:**
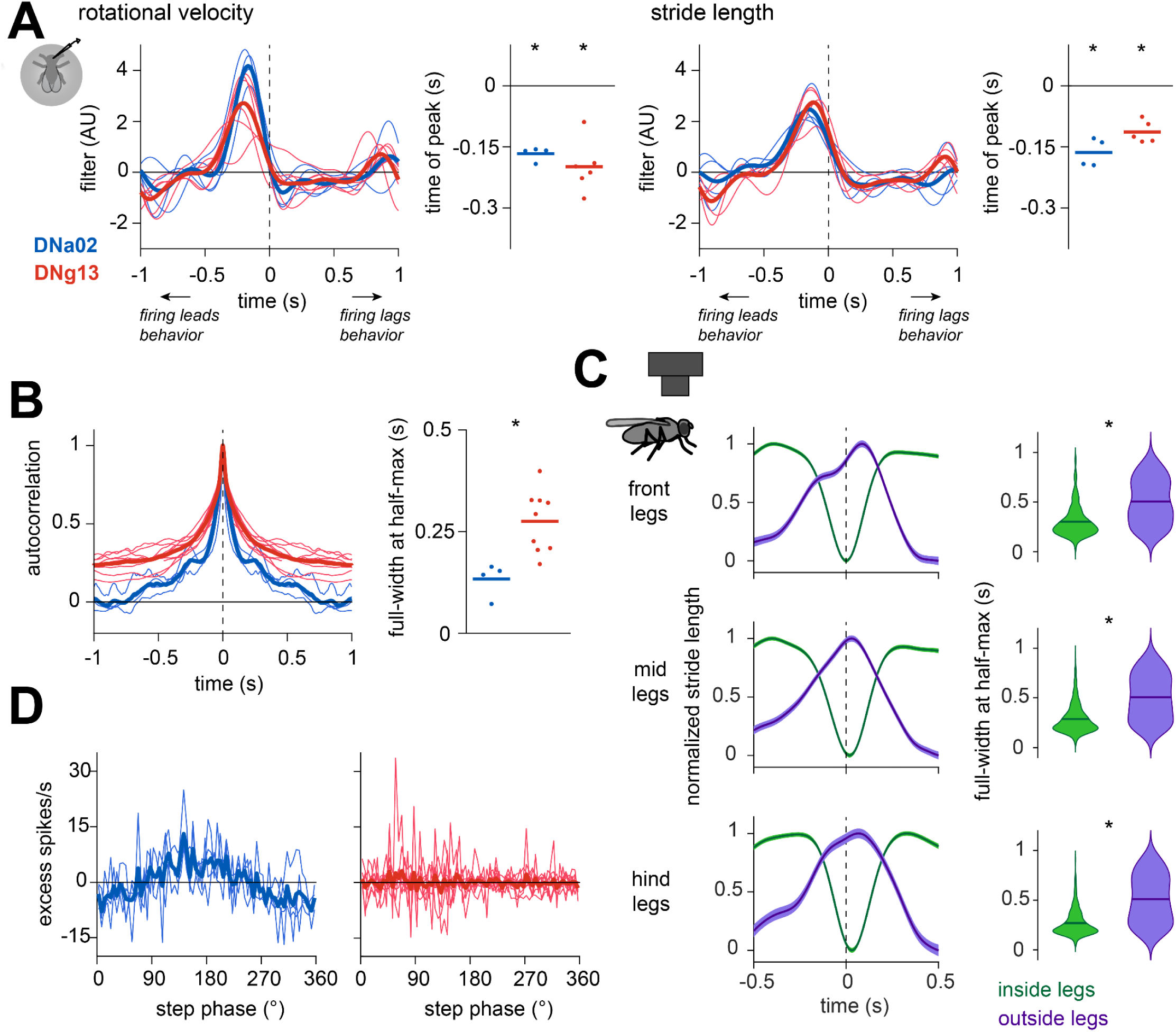
Different descending neurons are recruited at different moments within a steering maneuver. (A) *Left*: Filters describing the mean change in rotational velocity around an impulse change in DN firing rate. Thin lines are individual flies, and thick lines are the means across flies (n=4 cells in 4 flies for DNa02, 5 cells in 5 flies for DNg13). Stripchart shows time of the filter peak; dots are individual flies, and horizontal lines are the means across flies (one-sample t-test, time of peak different from 0, DNa02: p=2.85×10^−4^, DNg13: p=5.36×10^−4^). *Right*: Filters describing the mean change in stride length around an impulse change in DN firing rate. Stride length is measured in ipsilateral legs for DNa02 and contralateral legs for DNg13. For ease of comparison, the filter for DNa02 was multiplied by −1. Stripchart shows time of the filter peak (one-sample t-test, time of peak different from 0, DNa02: p=0.00250, DNg13: p=7.24×10^−4^). (B) Autocorrelation of DNa02 and DNg13 spike rate during walking (n=4 cells in 4 flies for DNa02, 9 cells in 9 flies for DNg13). Stripchart shows the width of the autocorrelation function (full-width at half-maximum). The two DNs are significantly different (two-sample t-test, p=0.00520). (C) *Left:* Changes in stride length over time for each pair of legs during turning bouts in freely walking flies. Turning bouts are aligned to the peak in rotational velocity (n=818 turning bouts in 73 flies). *Right*: The full width at half-maximum for the change in stride length over time. The duration of ipsilateral and contralateral changes in stride length are significantly different (paired t-test for each pair of legs, front: p=1.31×10^−76^; mid: p=8.08×10^−84^; hind: p=4.95×10^−93^). (D) Timing of DNa02 and DNg13 spikes relative to the stride cycle 100 ms prior (n=4 cells in 4 flies for DNa02, 9 cells in 9 flies for DNg13). The stride cycle is expressed relative to the phase of the ipsilateral (DNa02)/contralateral (DNg13) mid leg, and the binned spike counts are normalized to the time spent in each phase and then expressed as the difference from the mean.

Interestingly, DNg13 firing rate modulations were slower than DNa02 firing rate modulations, as evidenced by the different width of each cell’s autocorrelation function (Figure 6B). This difference is notable because it mirrors the way that ipsilateral and contralateral legs are coordinated during steering maneuvers. Specifically, during a turning bout, we found that stride length on the outside of the turn tends to slowly increase and decrease, while stride length on the inside of the turn changes more abruptly and transiently (Figure 6C). Our data suggest that the timing of this pattern originates in the brain, at the level of DN dynamics.

We wondered whether some of the fine structure in DNa02 spiking, which can be somewhat irregular when flies turn rapidly, was locked to the stride cycle. We therefore extracted the phase of the stride cycle at every time point in all our experiments, and we binned all DN spikes according to this phase. Indeed, we found that DNa02 spikes occurred preferentially at a specific phase of the stride cycle, whereas DNg13 spikes did not (Figure 6D). Although this stride-locked firing rate modulation was only about 15 spikes/sec, it was highly consistent across DNa02 recordings. Because the locomotor rhythm is intrinsic to the ventral nerve cord^3–7^, this finding suggests that DNa02 activity is influenced by ascending projections from the cord^67,68^.

### Descending neuron connectivity implies dynamic gating in the ventral nerve cord

Finally, we took advantage of the ventral nerve cord connectome^25^ to ask whether the inferred function of each DN is compatible with its synaptic targets in the cord (Figure S6A). Consistent with a recent analysis^69^, we found that both DNa02 and DNg13 synapse mainly onto premotor neurons–i.e. neurons presynaptic to motor neurons (Figures 7A and S6B); the DNs also make a few direct connections onto motor neurons. Each DN type distributes its output connections to all three pairs of legs (Figures 7B and S6C) but with essentially no overlap between the two DN types (Figures 7C and S6D).

**Figure 7:**
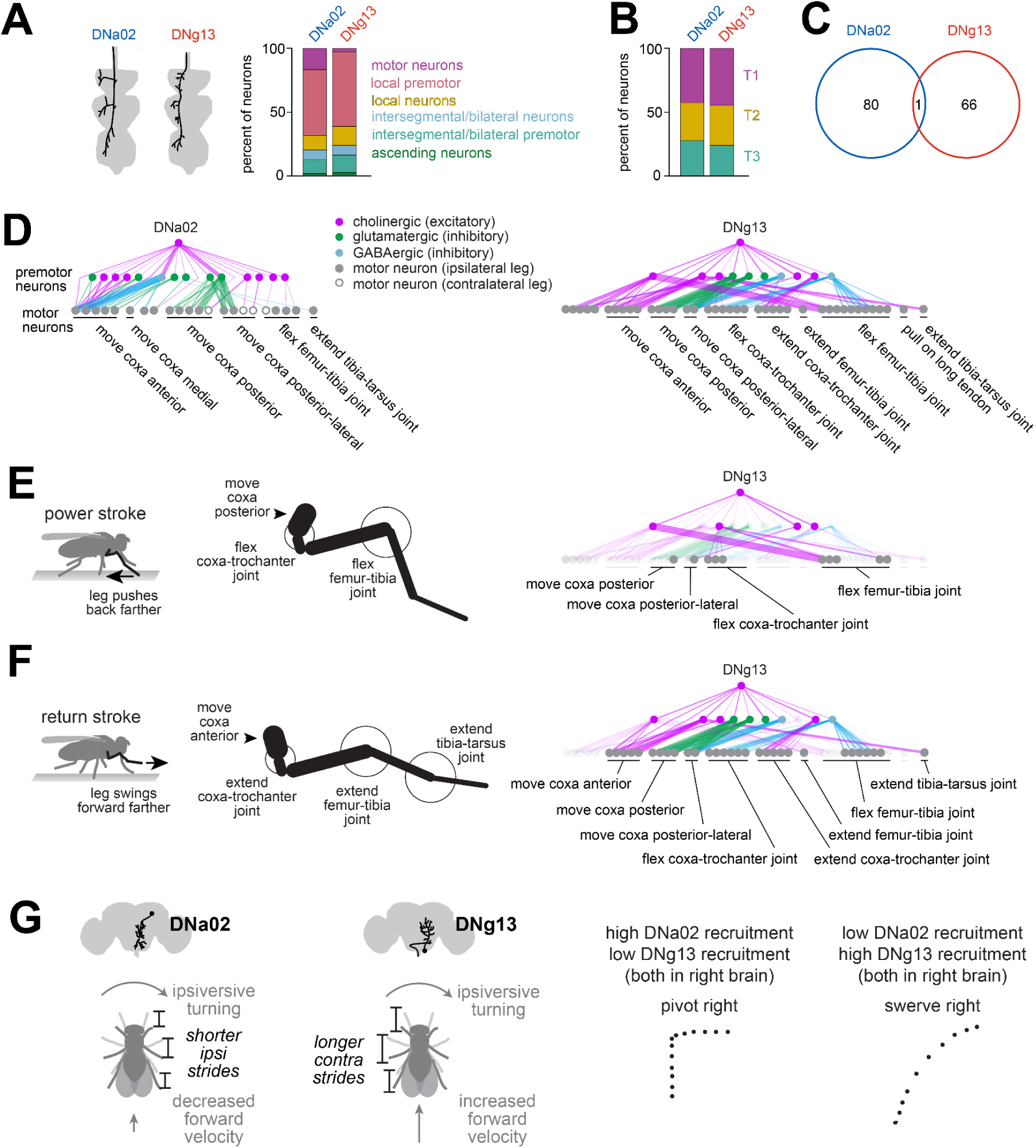
Steering descending neurons have specialized outputs. (A) Schematic: morphology of the left DNa02 and the right DNg13 axons in the ventral nerve cord. Here, left and right refer to soma locations in the brain; both of these cells send axons to the left side of the cord. Stacked bar charts summarize the cells postsynaptic to each DN in the leg neuromeres of the cord, divided into 6 categories. Note that DNa02 also targets neurons in the neuropils associated with the wings, halteres, and neck, and those are excluded from this plot. (B) The distribution across the three leg neuromeres (T1, T2, T3) of the postsynaptic neurons. Intersegmental neurons are assigned to a segment based on their cell body locations. (C) DNa02 and DNg13 axons projecting to the left side of the cord share almost no postsynaptic cells (numbers are cell counts). (D) Monosynaptic and disynaptic connections between DNa02/DNg13 axons projecting to the left side of the cord and motor neurons controlling the front legs. Note that the DNa02 and DNg13 graphs are plotted separately, and thus some motor neurons are repeated. Motor neurons are labeled according to their inferred effect^70^, with unlabeled motor neurons having unknown effects. Motor neurons are sorted in proximal-to-distal order of their target muscles. Neurotransmitter labels reflect machine vision predictions^72^ (for DNs) or hemilineage membership^71,103,104^ (for premotor cells). Line thickness is proportional to the number of synapses. Connections between premotor neurons are not shown. (E) *Left*: limb kinematics during the power stroke. *Right*: cells downstream from DNg13 that can potentially explain its ability to increase stride length during the power stroke. (F) Same but for return stroke. For DNa02 the relationship between motor neuron connections and limb kinematics is less clear. (G) Schematic summarizing the effects of each DN type on stride length, body velocity, and path shape.

To better understand the logic of these connections, we focused on the region of the ventral nerve cord that controls the front legs — the T1 neuromeres — because T1 motor neurons have been mapped to their muscle targets and actions on the leg joints^70^. In T1, we identified all the monosynaptic and disynaptic outputs of DNa02 and DNg13, and we determined the likely neurotransmitter for each disynaptic connection by identifying the hemilineage of the interposed cell, as neurons within the same hemilineage generally release the same neurotransmitter^71,72^.

We found that DNa02 and DNg13 are upstream from many motor neurons that collectively control muscles throughout the leg (Figure 7D). Specifically, DNg13 is positioned to increase the leg’s displacement during the power stroke and the return stroke, consistent with our functional data. During the power stroke, DNg13 is likely to excite motor neurons that move the coxa posterior, motor neurons that flex the femur-tibia joint, and motor neurons that flex the coxa-trochanter joint (Figures 7D and 7E). All these actions should lengthen the power stroke. Then, during the return stroke, DNg13 is positioned to excite motor neurons that move the coxa anterior, motor neurons that extend the coxa-trochanter joint, motor neurons that extend the femur-tibia joint, and motor neurons that extend the tibia-tarsus joint, while also inhibiting motor neurons that move the coxa posterior, motor neurons that flex the coxa-trochanter joint, and motor neurons that flex the femur-tibia joint (Figures 7D and 7F). All these actions should lengthen the return stroke. In short, connectome data are consistent with our finding that DNg13 increases stride length by modulating both phases of the step cycle. Importantly, however, this anatomy is only compatible with our results if we assume that incoming descending signals are dynamically gated by the locomotor cycle in the ventral nerve cord.

The situation for DNa02 is more complicated. During the return stroke, DNa02 is positioned to strongly inhibit motor neurons that move the coxa anterior (Figure 7D), which could shorten the return stroke, consistent with our functional data. That said, DNa02 is also positioned to excite these same motor neurons; it is also positioned to inhibit the motor neurons that move the coxa posterior (Figure 7D). Knowing how these signals are gated by the locomotor rhythm of the ventral nerve cord should help to clarify the logic of this connection pattern.

As an aside, we were surprised to discover that some cells postsynaptic to DNa02 were wing, haltere, or neck motor neurons or premotor neurons (Figures S6E and S6F). This was not the case for DNg13, whose direct postsynaptic neurons were all related to leg control (Figure S6F). This connectivity pattern suggests that DNa02 contributes to steering during flight as well as walking, whereas DNg13 is more specifically involved in steering during walking.

## Discussion

### Elemental components of locomotor control

The components of motor control at the level of the spinal cord or ventral nerve cord are not necessarily the same as the components of motor control from the brain’s perspective. In particular, during locomotion, the cord is responsible for generating the cyclical patterns of muscle activation that produce alternating power strokes and return strokes, via recruitment of muscle synergies^22^. Meanwhile, the brain is thought to control higher-level features of locomotion, such as heading direction. But there are also more granular components of locomotion that the brain might have access to, such as coordinated kinematic changes in specific combinations of limbs (gestures). To understand the descending control of locomotion, we need to identify what sorts of components are relevant to the brain.

Here we specifically focused on the elements of steering control in walking *Drosophila*. We found that different steering maneuvers involve different combinations of leg gestures. Specifically, one gesture (increasing stride length on the outside of a turn) was more prominent in swerves versus pivots, whereas another gesture (decreasing stride length on the inside of a turn) was more prominent in pivots versus swerves. Our perturbation results demonstrate that DNg13 can evoke the first gesture, while DNa02 can evoke the second gesture. Moreover, DNg13 and DNa02 are downstream from largely non-overlapping networks in the brain, implying that these DNs can be recruited independently. We propose that, by varying the ratio of activity in these two DNs, the brain can control the generation of pivots versus swerves (Figure 7G). Here we have focused on the simplest possible taxonomy of steering maneuvers, involving just two types of turns; in reality, there is a graded spectrum of steering maneuvers during walking^57^ which could arise from graded variations in the recruitment of DNg13 and DNa02 and likely other steering DNs as well^30,31,34,56^.

Strikingly, the function we have identified for DNa02 resembles that of a specific DN type in the mammalian brainstem, called Chx10. Unilateral activation of Chx10 cells reduces stride length on the ipsilateral side of the body, resulting in an ipsiversive turn, while also producing a transient decrease in forward velocity^20^. Our results expand upon that work by showing that, in *Drosophila*, DNa02 plays an analogous role, and this exists in parallel with another descending cell type that lengthens strides contralaterally (DNg13). Together, these two descending pathways can explain why it is possible for forward velocity to either increase or decrease during a turn (Figure 7G).

Our results support a model where the speed and direction of locomotion are each influenced by the activity of multiple DNs, with some DNs contributing to both^29–32,34,48,56^. Both parameters are also under feedback control, as sensory reafference continuously informs the brain about the body’s instantaneous forward velocity^73–75^ and heading^76^. Our results help illustrate how these feedback control loops can recruit specific DNs. For example, DNa02 is a major target of the central complex^31,61^, the brain region that compares the animal’s current heading to an internal goal direction^62,63,76,77^. When the central complex commands a large and abrupt change in heading, it also commands a sudden decrease in forward velocity^62,77^ — in other words, the central complex tends to drive pivoting maneuvers. This makes sense: if the brain’s course control system has just concluded that the animal has been moving in the wrong direction, then forward movement should be reduced until the error can be corrected. It is therefore notable that the leg gestures downstream from DNa02 activation are disproportionately associated with pivoting maneuvers, as we show here. Together, these results argue that the brain’s dedicated system for monitoring and controlling head direction is tightly coupled to the DNs that control pivoting.

Conversely, our data imply that DNg13 functions to change heading without decreasing forward velocity. Interestingly, this DN is not a direct target of the central complex; rather, it is downstream from other networks, including visuomotor circuits. This cell may be recruited by brain networks that stabilize straight walking, correcting for small, involuntary body rotations while continuing the body’s forward progression. This type of course control does not need to keep track of heading; rather, its function would simply be to stabilize straight-line locomotion, regardless of the direction of travel.

In the future, we expect that the *Drosophila* connectome will help us understand the functions of other DN types involved in steering^30,31,52,56^. For example, we predict that there exists a DN type that controls the stride length of the inside legs by modulating the power stroke, since DNa02 is specific to the return stroke but freely walking flies modulate both. Other DNs may control additional features of steering-related leg movements^31,36,37^ and body movements^57,78,79^. Finally, some DNs may only be recruited under specific internal states: for example, DNp09 has been suggested to control male steering during courtship^34^, but we observed this cell type was not active in our experiments (which focused on isolated females rather than courting males), consistent with previous work^34^.

### Rhythmic activity in descending neurons

Locomotor rhythms are prominent in the vertebrate brainstem^80–83^, cortex^84^, and cerebellum^85^. These rhythms are generally attributed to ascending projections from the spinal cord to the cerebellum and brainstem^9^, but vestibular signals may also contribute^86^. Interestingly, locomotor rhythms are prominent in descending cells that drive steering^83,87,88^, and it has been suggested that these rhythmic descending signals are timed to produce a laterally asymmetric modulation of rhythmic muscle contractions at appropriate times in the locomotor cycle^89^. However, we do not yet have a detailed understanding of how these descending cells orchestrate steering, and so the function of their rhythmicity is still uncertain.

Here we show that certain *Drosophila* steering DNs (DNa02 cells) are also phase-locked to the locomotor cycle. Because the *Drosophila* walking rhythm is intrinsic to the ventral cord^6^, phase-locking in DNa02 is almost certainly due to ascending projections^67,68^. Indeed, we found that DNa02 receives substantial monosynaptic input from ascending axons. Future work will be needed to determine whether the walking rhythm in DNa02 has a common origin with the walking rhythm reported in the *Drosophila* visual system^90^.

Interestingly, we also found that DNa02 specifically modulates one phase of the step cycle, the return phase. The return phase occurs at a different time in the front/back legs versus the middle legs^36,37,91^. We can thus infer that this single DN must have asynchronous effects on different legs. If the spikes traveling down this axon are phase-locked in order to arrive at the appropriate time, this means that the “appropriate time” must have a different phase relationship in different legs.

In summary, it seems that the timing of descending steering commands emerges from a combination of rhythmicity (due to ascending input) and limb-specific phase shifting. This is an economical architecture. Rather than sending a differently phase-shifted signal to different legs, the brain sends a single rhythmic signal to all the homolateral legs, and then relies on local mechanisms, such as proprioceptive feedback or the dynamics of the premotor circuits, to time-shift the effect of this signal.

### Opposing effects of single descending neurons

In vertebrates, stimulation of specific brainstem loci can evoke opposing effects on limb kinematics, depending on the phase of the locomotor rhythm when the stimulus arrives^17,92^. There are two possible explanations for this phenomenon. First, the same DN could have opposing effects on spinal targets, depending on the phase of the locomotor rhythm. Alternatively, the stimulus might recruit DNs with opposing effects, which are then gated at the level of the spinal cord in an alternating manner.

Here we show that, in *Drosophila*, a single DN can in fact have opposing effects, depending on the phase of the locomotor cycle. Specifically, DNg13 drives the legs to a more posterior position during the power stroke; conversely, it drives the legs to a more anterior position during the return stroke. In other words, it drives increased flexion in one phase, and increased extension in another phase. This finding implies that the downstream effects of this cell are inverted during the two phases of the step cycle.

Our analysis of DNg13 connectivity in the ventral nerve cord provides a mechanism to explain this phenomenon. We found that DNg13 is positioned to flex and extend multiple leg joints via distinct interposed interneurons in the ventral nerve cord. It seems likely that these interposed interneurons are rhythmically locked to the step cycle, so that descending drive is re-routed to these two effector pathways in an alternating manner. This would produce increases in stride length in both phases of the step cycle.

In short, our results demonstrate that a DN can have opposing effects, depending on the state of local pattern generators in the ventral cord. Again, this is an economical architecture because it minimizes the number of descending axons that must pass through the neck connective. Rather than devoting one DN to increasing flexion and another DN to increasing extension, the brain sends a common descending signal whose “meaning” is changed locally to produce both effects in an alternating sequence.

### Beyond arthropods

There is good reason to think that locomotor control in arthropods is relevant to vertebrates. First, arthropods and vertebrates share the same basic walking rhythm, with alternating power and return phases^2,93^. In both arthropods and vertebrates, motor neuron recruitment follows the same basic relationship between force production and recruitment order^94^. Moreover, arthropods and vertebrates confront shared physical problems, such as the need to balance stability and maneuverability^95,96^. Also, they confront common cognitive problems in complex locomotor control: arthropods navigate toward faraway destinations, memorized locations, or remembered objects^97,98^, just as vertebrates do. The links between navigation and locomotor control are unclear in vertebrates, but these links have recently been described in arthropods^62,63,75^. Given that brain circuits for navigation have clear homologies in arthropods and vertebrates^99,100^, it seems likely that insights from arthropods will have general relevance.

Our results provide further support for the idea of deep homology between locomotor control systems in arthropods and vertebrates. The functional similarity of DNa02 (in *Drosophila*) and Chx10 (in mammals^20^) is perhaps the most notable example of this. If we can understand the logic of descending locomotor control in any species, it will allow us to re-frame the problems of navigation and locomotor planning as concrete problems of generating specific dynamic activity patterns in particular brain cells^101,102^.

## Acknowledgements

Thomas Clandinin shared *UAS-CyRFP1* and *w+ norpA[P24]* flies. Barrett Pfeiffer and Gerald Rubin shared {*20XUAS-IVS-mCD8::GFP}attP40* flies. Kristin Branson and Alice Robie shared Animal Part Tracker software. John Tuthill and Anthony Azevedo shared cell type identifications in the FANC data. Greg Jefferis shared cell type identifications in the FAFB and FANC data. Jasper Phelps and Wei-Chung Lee provided access to FANC data. Gwyneth Card, Michael Dickinson, and Shigehiro Namiki shared DN split-GAL4 lines and confocal images. We thank the FlyWire and FANC communities for proofreading EM segmentation. This work was supported by a Jane Coffin Childs Fellowship (to H.H.Y.), an NIH K99 award (NS129759 to H.H.Y.), an NSF GRFP fellowship (to L.E.B.), an NIH R01 (NS070644 to R.S.M.), and a U19 team grant (NS104655 to R.I.W., R.S.M., Thomas Clandinin, Michael Dickinson, and John Tuthill). R.I.W. is an HHMI Investigator.

## Author contributions

H.H.Y. performed calcium imaging, optogenetics, and electrophysiology experiments. L.E.B. collected behavioral data from freely walking flies. L.S., Q.X.V., and A.A. annotated and proofread EM segmentation. H.H.Y. designed and performed data analyses. S.R. provided useful suggestions and shared analyses of unpublished data. R.S.M. provided supervision and funding for A.A., as well as useful suggestions. H.H.Y. and R.I.W. conceptualized the study and wrote the manuscript.

## STAR Methods

### Key resources table

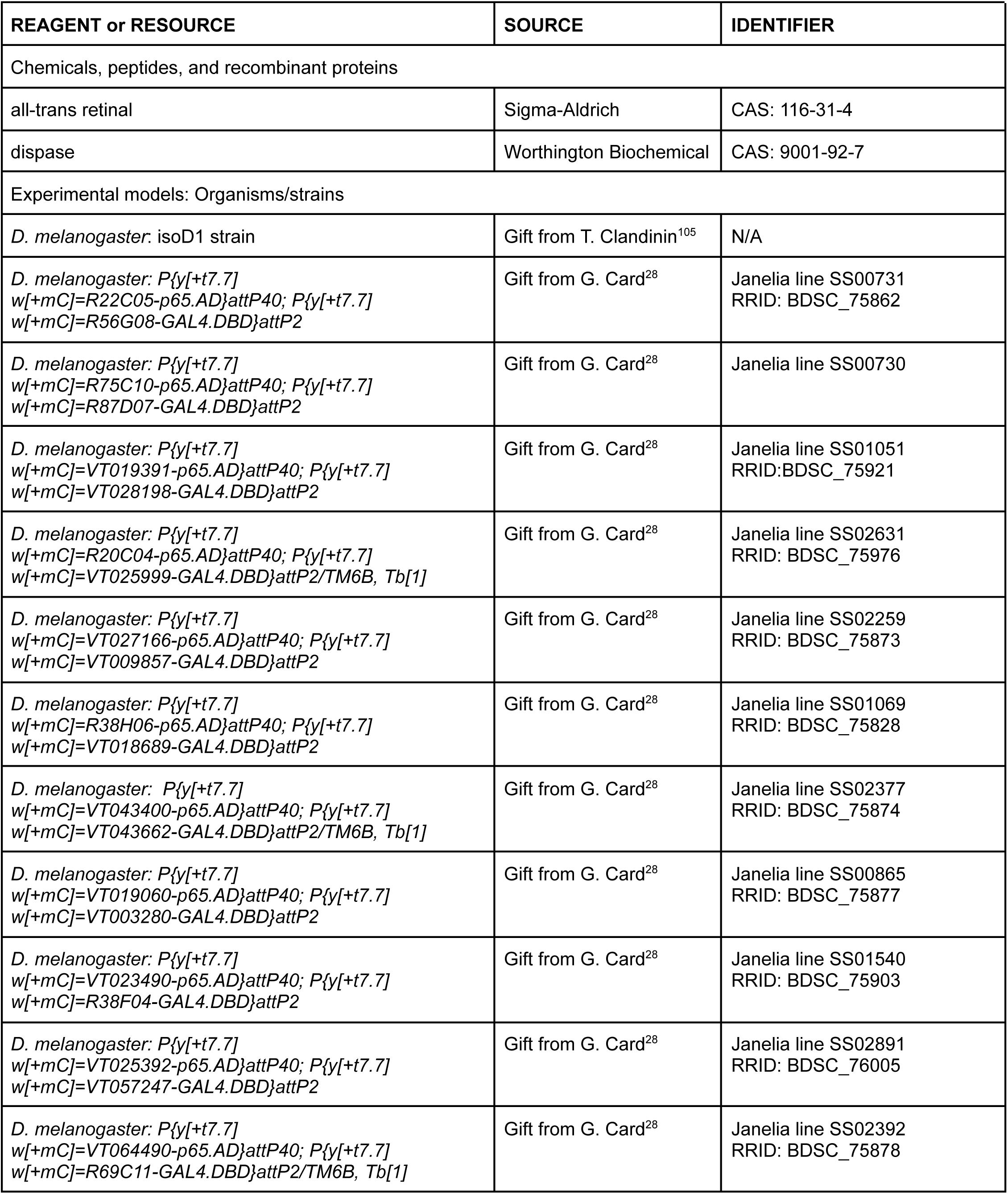
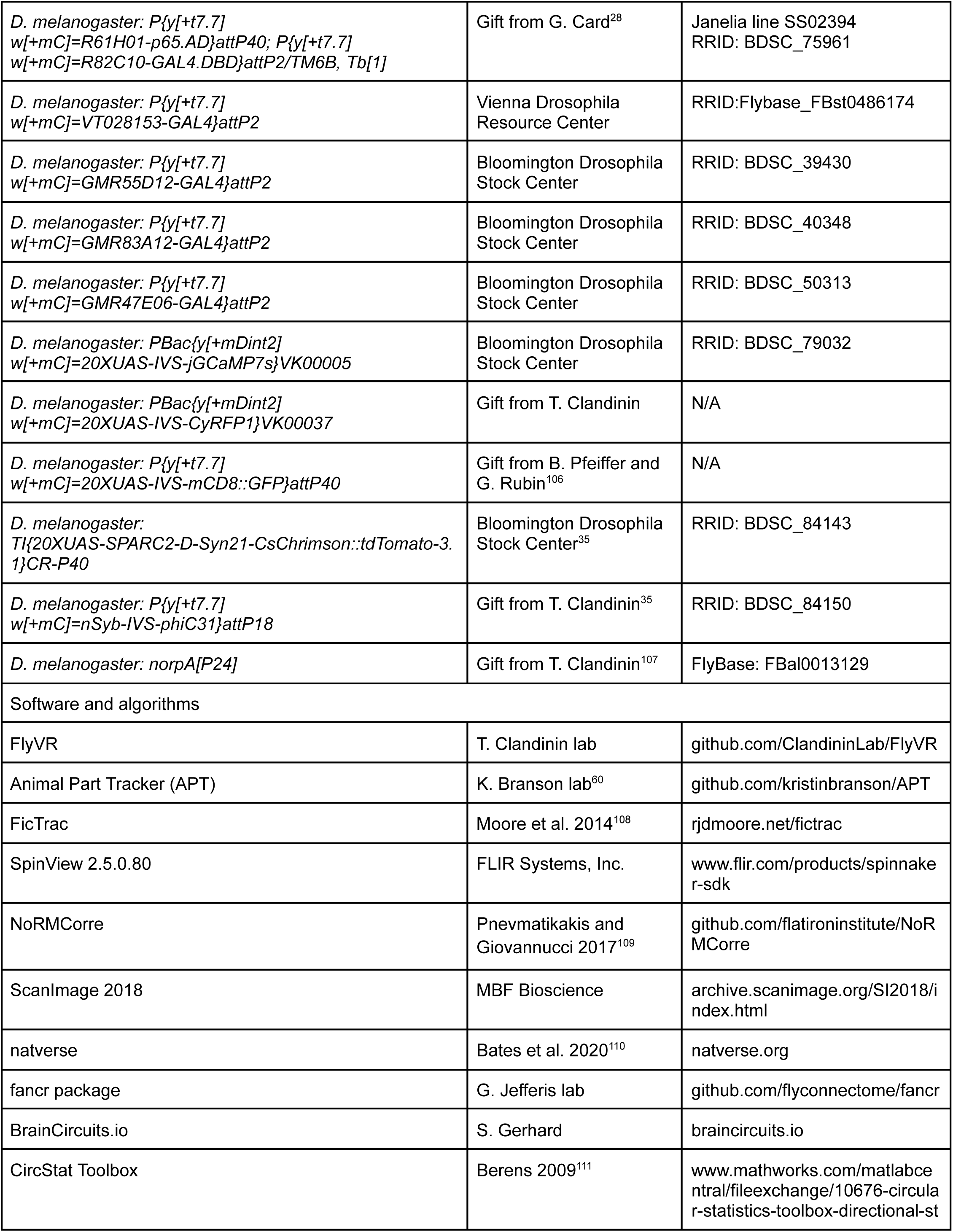
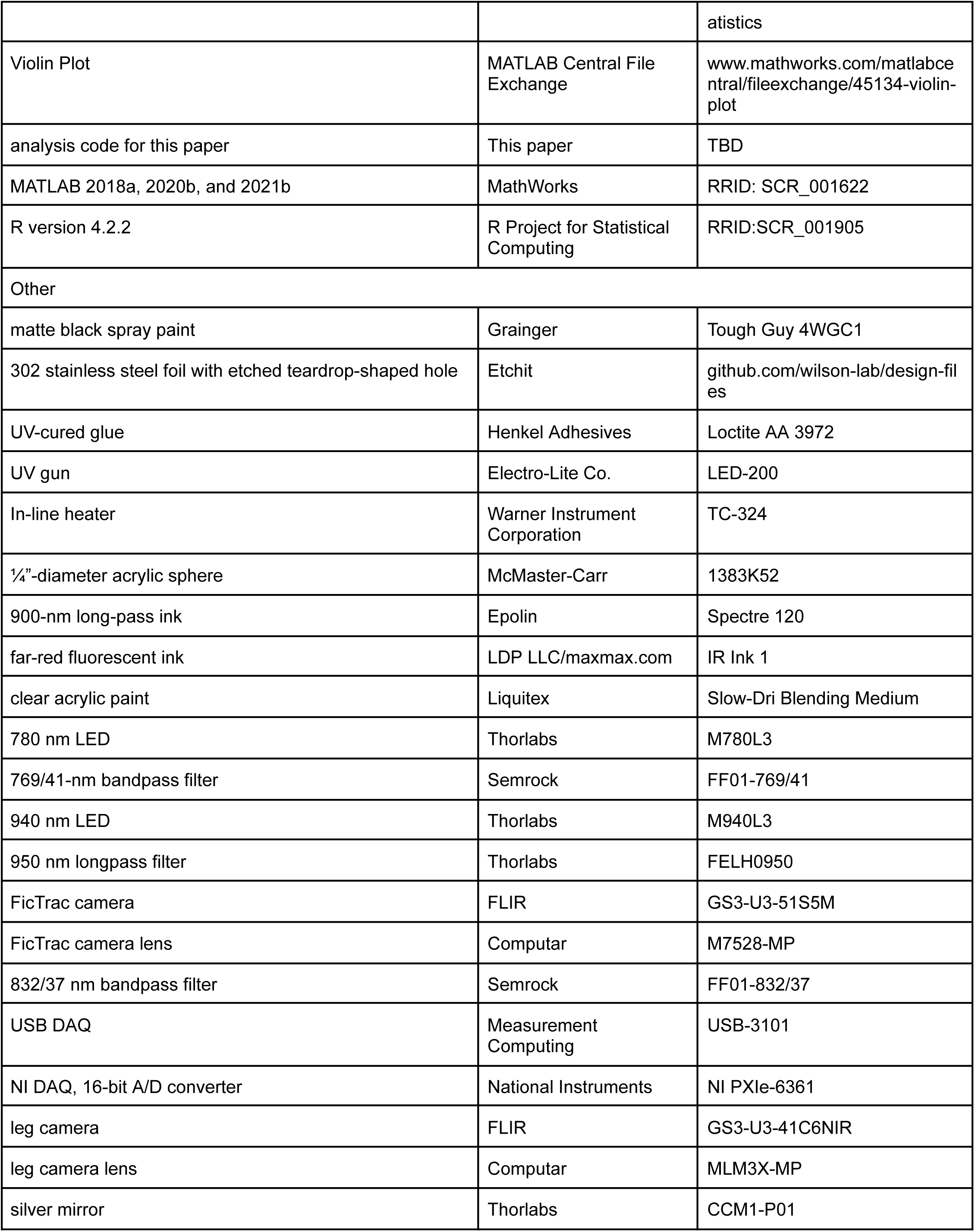
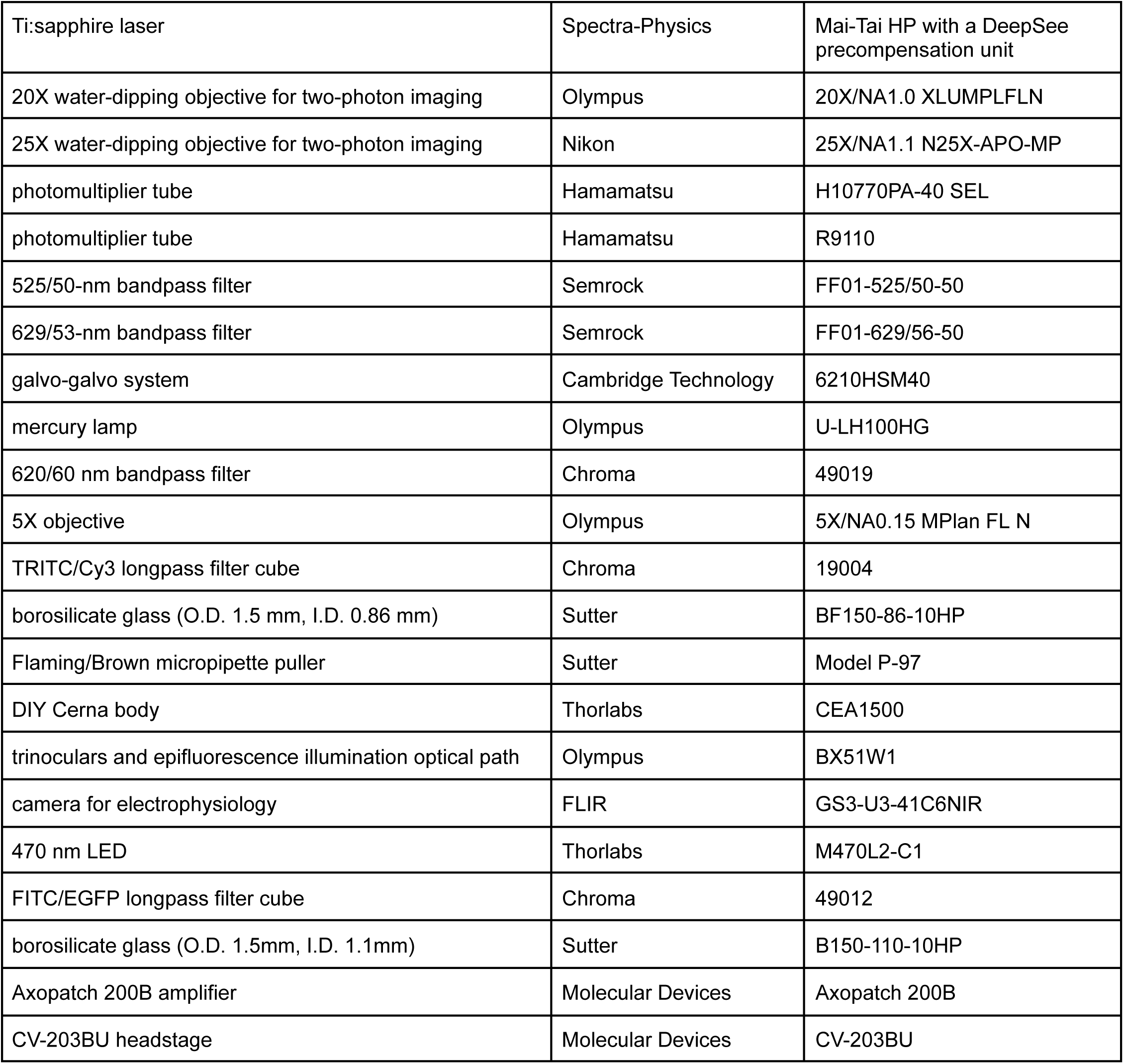

### Resource availability

#### Lead contact

Further information and requests for resources and reagents should be directed to and will be fulfilled by the lead contact, Rachel Wilson.

#### Materials availability

All reagents generated in this study are freely available upon request.

#### Data and code availability

All data reported in this paper will be shared by the lead contact upon request. All original code will be deposited at Zenodo and made publicly available. DOIs will be included in the key resources table. Any additional information required to reanalyze the data reported in this paper is available from the lead contact upon request.

### Experimental model and study participant details

#### Drosophila melanogaster

The isoD1 strain of *D. melanogaster* was used as wild-type^105^. All transgenes, strains, and mutant alleles used in this study are listed in the Key resources table. All experimental flies were female. All flies except those used for optogenetic stimulation experiments were raised on standard cornmeal-molasses food (recipe B2, Archon Scientific) at 25 °C, 50-70% humidity, and a 12:12 light:dark cycle. Flies used for optogenetic stimulation experiments were raised at 25 °C and 50-70% humidity on semi-defined food^112^, following the BDSC protocol, supplemented with all-trans retinal (ATR, Sigma-Aldrich) at a final concentration of 0.6 mM. Because ATR is light-sensitive, vials were individually wrapped in foil, and the flies were raised in constant darkness.

For freely walking fly behavior experiments, female flies were collected on CO_2_ and housed in groups of 10-20 on cornmeal-molasses food until they were used for the experiments at 2-5 days post-eclosion. On the day of the experiment, individual vials were loaded into the automated dispenser. Experiments were performed during the flies’ subjective morning (within ∼3 hours of the dark to light transition, Zeitgeber time 0).

For tethered fly experiments, female flies were anesthetized on ice for collection within 24 hours of eclosion. For calcium imaging experiments, most genotypes were used 1-3 days post-eclosion, but DNp05 flies were used 10-16 hours post-eclosion as the expression of jGCaMP7s was higher in younger flies. Flies were wet-starved by housing them in a vial with a water-dampened kimwipe but no food for 20-30 hours prior to the experiment to encourage walking. For flies used at 1 day or less post-eclosion, this meant that they were starved from collection until experiment. For flies used at older ages, they were housed on cornmeal-molasses food upon collection until the starvation period. Experiments were performed during the flies’ subjective evening (± 3 hours from the light to dark transition, Zeitgeber time 12). For optogenetic experiments, flies were collected onto semi-defined food with 0.6 mM ATR and housed in constant darkness. They were used for experiments at 7 days post-eclosion to allow time for the SPARC reaction to occur and CsChrimson to be expressed. For the 20-30 hours prior to the experiment, these flies were wet-starved by housing them in a vial with a kimwipe dampened with water and 1mM ATR. For electrophysiology experiments, flies were used 1 day post-eclosion. Like the calcium imaging flies, they were wet-starved for 20-30 hours post-eclosion, and experiments were performed in the flies’ subjective evening.

### Method details

#### Fly genotypes by figure

Figures 1, S1, and 6C:

*+; +; +* isoD1 strain

Figure 2:

**DNa01**: w[1118]/+; R22C05AD/UAS-CyRFP1; R56G08DBD/UAS-jGCaMP7s
**DNa02:** *w[1118]/+; R75C10AD/UAS-CyRFP1; R87D07DBD/UAS-jGCaMP7s*
**DNb05:** *w[1118]/+; VT019391AD/UAS-CyRFP1; VT028198DBD/UAS-jGCaMP7s*
**DNb06:** *w[1118]/+; R20C04AD/UAS-CyRFP1; VT025999DBD/UAS-jGCaMP7s*
**DNg13:** *w[1118]/+; VT027166AD/UAS-CyRFP1; VT009857DBD/UAS-jGCaMP7s*

Figure S2:

**DNa01**: w[1118]/+; R22C05AD/UAS-CyRFP1; R56G08DBD/UAS-jGCaMP7s
**DNa02:** *w[1118]/+; R75C10AD/UAS-CyRFP1; R87D07DBD/UAS-jGCaMP7s*
**DNb05:** *w[1118]/+; VT019391AD/UAS-CyRFP1; VT028198DBD/UAS-jGCaMP7s*
**DNb06:** *w[1118]/+; R20C04AD/UAS-CyRFP1; VT025999DBD/UAS-jGCaMP7s*
**DNg13:** *w[1118]/+; VT027166AD/UAS-CyRFP1; VT009857DBD/UAS-jGCaMP7s*
**DNg14:** *w[1118]/+; R38H06AD/UAS-CyRFP1; VT018689DBD/UAS-jGCaMP7s*
**DNg15:** *w[1118]/+; VT043400AD/UAS-CyRFP1; VT043662DBD/UAS-jGCaMP7s*
**DNg16:** *w[1118]/+; UAS-CyRFP1/+; VT028153-GAL4/UAS-jGCaMP7s*
**DNg31:** *w[1118]/+; UAS-CyRFP1/+; GMR55D12-GAL4/UAS-jGCaMP7s*
**DNg34:** *w[1118]/+; UAS-CyRFP1/+; GMR83A12-GAL4/UAS-jGCaMP7s*
**DNp05:** *w[1118]/+; VT019060AD/UAS-CyRFP1; VT003280DBD/UAS-jGCaMP7s*
**DNp09:** *w[1118]/+; VT023490AD/UAS-CyRFP1; R38F04DBD/UAS-jGCaMP7s*
**DNp11:** *w[1118]/+; VT025392AD/UAS-CyRFP1; VT057247DBD/UAS-jGCaMP7s*
**DNp12:** *w[1118]/+; UAS-CyRFP1/+; GMR47E06-GAL4/UAS-jGCaMP7s*
**DNp18:** *w[1118]/+; VT064490AD/UAS-CyRFP1; R69C11DBD/UAS-jGCaMP7s*
**DNp32:** *w[1118]/+; R61H01AD/UAS-CyRFP1; R82C10DBD/UAS-jGCaMP7s*

Figures 3D, 4A-C and S4 (DNa02 optogenetic stimulation):

norpA[P24], nSyb-PhiC31/norpA[P24]; UAS-SPARC2-D-CsChrimson::tdTomato/R75C10AD; R87D07DBD/+

Figures 3D, 4D-F, S4D (DNg13 current injection) and Figures 5A, 5B, 5D, 5G, S5A, S5C, S5D, 6A, 6B, and 6D (DNg13 electrophysiology recordings):

*+; UAS-mCD8::GFP/VT027166AD; VT009857DBD/+* (backcrossed 5 generations into the isoD1 genetic background)

Figure 5A, 5B, 5C, 5E, 5F, S5A, S5B, S5D, 6A, 6B, and 6D (DNa02 electrophysiology recordings):

*+; UAS-mCD8::GFP/R75C10AD; R87D07DBD/+* (backcrossed 5 generations into the isoD1 genetic background)

#### Freely walking fly behavior

The behavior of freely walking flies was collected using the Coliseum^59^. Briefly, the Coliseum consisted of a 1m x 1m arena with a large monitor forming each of the 4 walls. Flies were automatically and individually dispensed into the arena through a hole in the floor using an automated dispenser. The flies were illuminated from below the floor using IR LEDs and tracked from above using a high-speed camera mounted on a 2-axis CNC (grbl with gShield + Arduino UNO; stepper motors SM42HT47–1684B), which maintained the fly in the camera’s field of view in closed-loop using the FlyVR software (github.com/ClandininLab/FlyVR). The video of the fly as well as the fly’s body position and angle in the frame and the position of the CNC were output by FlyVR at the end of each trial. Trials ended when tracking was lost for several seconds (because the fly left the arena by flying away). The frame rate of position tracking and the video was ∼125 Hz. Video frames were 658 px x 494 px, and the resolution was 37 px/mm. Flies walked with a constant background (dark, gray, or bright) or a static pattern (vertical or horizontal sine gratings or a checkerboard) on the monitor walls around them; for subsequent analyses, flies across all of these wall backgrounds/patterns were pooled.

#### Tethered fly preparation and dissection

The fly preparation holder consisted of a 302 stainless steel foil with a bend in it secured to an acrylic platform. The bottom side of the holder was spray-painted a solid matte black (4WGC1, Tough Guy) to reduce reflections during behavioral monitoring (see Tethered fly behavioral setup). The foil had a teardrop-shaped hole photochemically etched into it (Etchit), into which the fly’s head and thorax were gently pushed. The fly was then secured to the foil with UV-cured glue (Loctite AA 3972) that was cured with brief (less than 1 s) pulses of UV light from a UV gun (LED-200, Electro-Lite Co.). For calcium imaging and electrophysiology experiments, the fly’s head was angled to allow access to the desired arbors or cell bodies. For calcium imaging, this meant that for most genotypes, the fly’s head was tilted forward such that the posterior cuticle was exposed; however, for DNg14, DNg15, DNg16, and DNg31, the head was gently rotated 180° around the neck such that the ventral side of the head was facing upwards. For optogenetic experiments, the fly’s head was secured at approximately its natural angle. For DNa02 electrophysiology experiments, the fly’s head was tilted backwards until it hit the thorax to expose more of the anterior surface, though the antenna were kept below the foil. For DNg13 electrophysiology experiments, the head was rotated 180° around the neck to expose the ventral side. The bend in the foil allowed the angle between the thorax and the abdomen to be better matched to that of freely walking flies, which was important for achieving good walking in these tethered flies. To prevent excessive grooming of the wings and for easier automated tracking of the legs, the blades of the wings were fully removed, but the wing hinges were left attached.

For calcium imaging and electrophysiology experiments, the dorsal side of the fly was covered in saline, and to expose the brain area of interest, forceps were used to make a hole in the head capsule and to remove overlying trachea and fat. To reduce movement of the brain, muscle 16 was severed with a hook fashioned out of tungsten wire. Additionally, muscles of the proboscis were severed using forceps, or in the case of flies with their head rotated 180°, the proboscis was entirely removed to provide access to the brain. For electrophysiology experiments, a hole was made in the perineural sheath in the area around the cell bodies of interest with forceps. For optogenetic experiments, the head capsule was not opened and the proboscis was left intact. However, as described below, saline was still perfused over the dorsal side of the fly to control the temperature and to buffer the thermal effects of the optogenetic stimulation.

The external saline solution contained (in mM): 103 NaCl, 3 KCl, 5 *N*-tris(hydroxymethyl) methyl-2-aminoethane-sulfonic acid, 8 trehalose, 10 glucose, 26 NaHCO_3_, 1 NaH_2_PO_4_, 1.5 CaCl_2_, and 4 MgCl_2_, with the osmolarity adjusted to 270–273 mOsm. The saline was bubbled with 95% O_2_ and 5% CO_2_, and it equilibrated at a pH of 7.3. The temperature was maintained at 28-30°C with an in-line heater (TC-324, Warner Instrument Corporation). The external saline was continuously superfused at a rate of approximately 2 mL/min.

#### Tethered fly behavioral setup

For all calcium imaging, optogenetic stimulation, and electrophysiology experiments, the fly’s walking behavior was monitored using a custom-built setup that enabled simultaneous extraction of the fly’s intended walking trajectory as well as its leg movements. The fly walked on a ¼”-diameter acrylic sphere (1383K52, McMaster-Carr). This ball was first uniformly coated with 900-nm long-pass ink (Spectre 120, Epolin, diluted ∼2:3 in cyclohexanone), and then spots of far-red fluorescent ink (excitation 793 nm, emission 840 nm; IR Ink 1, LDP LLC/maxmax.com, mixed 1:1 with clear acrylic paint (Slow-Dri Blending Medium, Liquitex)) were painted on by blending into a still-wet coat of clear acrylic paint. For the calcium imaging experiments, the ball floated above a plenum made of a 0.6 mL microcentrifuge tube into which another ¼”-diameter acrylic sphere was inserted such that the distance between the surface of the two balls was ∼1.7 mm, which was the distance necessary for light passing through the two balls to be collimated. This 0.6 mL microcentrifuge tube was then inserted in a 1.5 mL microcentrifuge tube. Air was flowed into the 1.5 mL tube from two sides, and small holes were made in the wall of the 0.6 mL tube above the fixed ball to allow the air to flow to support the ball on which the fly walked. For the optogenetic and electrophysiology experiments, this microcentrifuge tube-based plenum was replaced with a 3D printed one of an equivalent overall design. The ball was illuminated with a 780 nm LED (M780L3, Thorlabs) that had a 769/41-nm bandpass filter (FF01-769/41, Semrock) in front of it. The fly was illuminated with two 940 nm LEDs (M940L3, Thorlabs) that each had a 950 nm longpass filter (FELH0950, Thorlabs) in front. The movement of the ball was tracked at ∼120 Hz in real time using FicTrac^108^, a program that computes the rotation of a sphere from video captured directly from a camera (here: GS3-U3-51S5M, FLIR; with lens M7528-MP, Computar; with a 832/37 nm bandpass filter, FF01-832/37, Semrock). FicTrac was modified to send real-time analog measurements of the position of the ball in all three rotational axes to a USB DAQ (USB-3101, Measurement Computing). These signals were digitized at 20kHz by a 16-bit A/D converter (NI PXIe-6361, National Instruments) and acquired using the MATLAB Data Acquisition Toolbox (R2018a, MathWorks). A video of the fly’s leg movements was captured from below at a resolution 69 px/mm (image size: 416 px x 416 px) at 223 Hz with a 4 ms exposure time (calcium imaging experiments) or at 250 Hz with a 1.5 ms exposure time (optogenetic stimulation and electrophysiology experiments) using a GS3-U3-41C6NIR (FLIR) camera with a macro zoom lens (MLM3X-MP, Computar) through the SpinView software (FLIR). The image was reflected off of a silver mirror (CCM1-P01, Thorlabs), and one or two 950 nm longpass filters (FELH0950, Thorlabs) were positioned between the bottom of the ball plenum and the camera. Two filters were required to exclude the light from the two-photon laser during calcium imaging experiments; one filter was sufficient to exclude stray light for optogenetic stimulation and electrophysiology experiments.

#### Two-photon calcium imaging

Calcium imaging experiments were performed on a custom-built two-photon microscope controlled with ScanImage software^113^. jGCaMP7s and CyRFP1 were excited at 920 nm using a Mai-Tai HP Ti:sapphire laser with a DeepSee precompensation unit (Spectra-Physics). Power at the sample was kept to under ∼20 mW. The objective used was either the 20X/NA1.0 XLUMPLFLN objective (Olympus) or the 25X/NA1.1 N25X-APO-MP objective (Nikon). jGCaMP7s fluorescence was collected using the H10770PA-40 SEL PMT (Hamamatsu) with a 525/50-nm bandpass filter (FF01-525/50-50, Semrock), and CyRFP1 fluorescence was collected using the R9110 PMT (Hamamatsu) with a 629/53-nm bandpass filter (FF01-629/56-50, Semrock). For all but DNp12, the imaging region for each cell type was a single plane selected to contain a portion of the dendritic arbor for both the left and the right cells. The imaging region for DNp12 instead contained the left and right cell bodies. The imaging region was 200 px by 80 px, and it was scanned bidirectionally using a galvo-galvo system (6210HSM40, Cambridge Technology) with a pixel dwell time of 2400 ns. The final frame rate was 22.7 Hz.

#### Optogenetic experiments

Illumination for stimulating CsChrimson was provided by light from a mercury lamp (U-LH100HG, Olympus) passed through a 620/60 nm bandpass filter (49019 - ET - Cy5 Longpass, Chroma) and focused onto the dorsal surface of the fly’s head and thorax using a 5X/NA0.15 MPlan FL N (Olympus) objective. The spot of illumination was 5.5 mm in diameter, and therefore covered the entire dorsal area of the fly exposed above the holder. ND filters were used to control the illumination intensity. For the data in Figures 3, 4, and S4, the illumination power delivered to the fly was 0.09 mW/mm^2^. For each fly, two 250-s trials were performed, and each trial consisted of randomly interleaved 0.2 s and 2 s light pulses separated by 5 s without stimulation.

Using SPARC, the experimental and control flies were of the same genotype, so the behavioral measurements were performed blinded to which DNa02 cells, if any, expressed CsChrimson. Afterwards, the brain and ventral nerve cord were dissected out of the fly and placed on a microscope slide. The expression of CsChrimson in each DNa02 cell was determined by the fluorescence of the tdTomato tagged to the CsChrimson under epifluorescence imaging (excitation filter: 540/25 nm, emission filter: 575 nm longpass, 19004 - AT - TRITC/Cy3 Longpass, Chroma). Expression of CsChrimson in non-DNa02 cells was also noted. In the brain, it was restricted to a single type of visual neuron with arbors in the lobula; on average, 2 cells per hemisphere expressed CsChrimson (range 0-4 cells). In the ventral nerve cord, a few cells in the abdominal ganglion expressed CsChrimson in a subset of flies.

SPARC-CsChrimson could not be used to produce specific unilateral stimulation of DNg13 because the transgenic line for this DN also drives GAL4 expression in an ascending neuron. In flies where this ascending neuron expressed CsChrimson, regardless of whether DNg13 expressed CsChrimson, light stopped locomotion. Very few flies had expression in one copy of DNg13 but zero copies of this ascending neuron, and so it was challenging to collect enough data to assess the effect of unilateral DNg13 stimulation alone. For this reason, direct current injection via patch-clamp electrophysiology was used to perturb DNg13 unilaterally.

#### Patch-clamp electrophysiology

Patch pipettes of borosilicate glass (O.D. 1.5 mm, I.D. 0.86 mm, Sutter) were made the day of the recording using a Flaming/Brown micropipette puller (Model P-97, Sutter). Pipette resistances were 4.5-6.5 MΩ. The internal solution contained (in mM): 140 potassium aspartate, 10 4-(2-hydroxyethyl)-1-piperazineethanesulfonic acid, 4 MgATP, 0.5 Na_3_GTP, 1 ethylene glycol tetraacetic acid, 1 KCl, and 15 neurobiotin citrate. The pH was 7.2 and the osmolarity was adjusted to 265 mOsm.

Patch-clamp recordings were achieved under visual control using a custom-built microscope made from a DIY Cerna body (Thorlabs) and the trinoculars and epifluorescence illumination optical path of an Olympus BX51W1. The camera was a GS3-U3-41C6NIR (FLIR). The objective was the 40x/NA0.8 LUMPlanFL N water dipping objective from Olympus. The brain was illuminated with the same light source as the ball (described above in *Tethered fly behavioral setup*); this was a 780 nm LED (M780L3, Thorlabs) that had a 769/41-nm bandpass filter (FF01-769/41, Semrock) in front of it. The desired cell was identified based on GFP expression and the location of the cell body. GFP was excited using illumination from either a 470 nm LED (M470L2-C1, Thorlabs) or a mercury lamp (U-LH100HG, Olympus) passed through a 480/40-nm filter (49012-ET-FITC/EGFP longpass, Chroma). The emission filter was a 510 nm longpass filter.

To expose the targeted cell body to access by the patch pipette, neighboring cell bodies and glia were first cleared away by applying positive and negative pressure through large-bore cleaning pipettes (pulled from O.D. 1.5mm, I.D. 1.1mm borosilicate glass, Sutter) filled with external saline. Ensheathing glia around DNg13’s cell body were removed by brief application of dispase (1mg/mL in external saline, Worthington Biochemical) delivered through a pipette.

Recordings were obtained using an Axopatch 200B amplifier and a CV-203BU headstage (Molecular Devices). Analog voltage signals were lowpass filtered at 5kHz (Bessel filter, 80 dB/decade), digitized at 20kHz by a 16-bit A/D converter (NI PXIe-6361, National Instruments), and acquired using the MATLAB Data Acquisition Toolbox (R2018a, MathWorks). Liquid junction potential correction was performed post-hoc by subtracting 13 mV from the recorded voltages^114^.

Hyperpolarizing current injection into DNg13 could suppress spike generation in this cell almost completely, whereas depolarizing current injection into DNg13 could reliably elevate the spike rate to >100 spikes/s. However, using current injection into DNa02, large and reliable changes in spike rate could not be generated. This could occur if the spike initiation zone in DNa02 is electrotonically distant from the soma. In any event, this precluded us from using current injection to create bidirectional perturbations of DNa02 activity, as we did for DNg13.

#### Data analysis - Freely walking fly behavior

Using Animal Part Tracker (APT)^60^ running the Mixture Density Model (MDN) deep convolutional network, the tips of the 6 tarsi, the front of the head, 3 points on the thorax (both postpronotal lobes and the posterior tip of the scutellum), the tip of the abdomen, and the 2 wing tips were tracked in each frame of the videos acquired from the Coliseum^59^. Using this position within the camera frame and the position of the CNC, the absolute position in world coordinates of each tracked point was extracted. From the tracked points, the position of the tarsi was extracted in units of body length in the coordinate frame defined by the fly’s anterior-posterior and medial-lateral axes. Specifically, a line was fit to the head, the midpoint between the postpronotal lobes, the posterior tip of the scutellum, and the tip of the abdomen to define the anterior-posterior axis of the body. The head, scutellum, and abdomen tracked points were projected to this axis; the distance between the head and abdomen projected points was defined as the fly’s body length and the scutellum point as the midpoint of the fly. The medial-lateral axis was defined as the axis orthogonal to the anterior-posterior axis that passed through the fly’s midpoint. The angle of the vector from head to abdomen along the anterior-posterior axis was the fly’s heading angle. This angle was used to disambiguate the heading angles computed by FlyVR (described below) but was not used to compute any body velocity parameters. After projecting the tarsi points to the fly’s coordinate frame, outliers were removed using the following procedure: the median filtered position was computed over a 5-frame window, and outliers that deviated by more than 50% of the range in median-filtered values over a 500-frame window were replaced with the median-filtered value. The maxima and minima of position in the anterior-posterior axis were extracted for each leg. Individual steps of the walking stride cycle defined as a single complete cycle from posterior to anterior to posterior, and the following step parameters were computed on each step:

- Anterior extreme position (AEP) - the position in both axes of the tarsal tip when it was at its most anterior position of the stride cycle
- Posterior extreme position (PEP) - the position in both axes of the tarsal tip when it was at its most posterior position of the stride cycle
- Stride length - the distance between the AEP and the PEP in the anterior-posterior axis
- Step direction - the angle of the vector from the AEP to the PEP
- Stance or swing - for each half step (posterior to anterior or anterior to posterior), the translational velocity of the leg in the world coordinate frame was computed. The half step where the translational velocity was smaller (i.e. the leg was stationary relative to the ground) was called stance, and the half step where the velocity was greater (i.e. the leg was moving relative to the ground) was called swing.
- Stance duration - the amount of time the leg spent in the stance phase
- Swing duration - the amount of time the leg spent in the swing phase
- Return stroke position - the AEP in the anterior-posterior axis at the end of the swing phase (i.e. only when the leg is walking forward)
- Power stroke position - the PEP in the anterior-posterior axis at the end of the stance phase (i.e. only when the leg is walking forward)

The phase of each leg during each frame was computed by assigning the phase to be 0° for the frame at the start of the swing phase for each step (based on the swing/stance calls), 180° for the frame at the swing/stance transition, and 360° for the frame at the end of the stance phase, and then for each half step, linearly interpolating between these values using the position in the anterior-posterior axis. Spline fits to the maxima and minima of position in the anterior-posterior axis were used to compute continuous estimates of the AEP and PEP. The difference between the AEP and PEP was the continuous estimate of stride length. And the angle of the vector from the AEP to the PEP was the continuous estimate of step direction.

The FlyVR software extracted the position and angle of the fly within the camera frame by fitting an ellipse to the body; the body position was the centroid of the ellipse, and the body angle was the angle of the long axis of the ellipse. This measure of body angle did not distinguish between the front and back of the fly, so it was compared with the fly’s heading angle from the leg tracking procedure, and any ∼180° deviations were corrected. The absolute body position in world coordinates was computed by summing the fly’s position in the camera frame with the position of the CNC. The body position and the heading angle were smoothed using a 10-frame gaussian-weighted moving average. The fly’s rotational velocity was the change in heading angle. The fly’s forward velocity was the change in position in the same direction as the heading angle. The fly’s lateral velocity was the change in position in the direction orthogonal to the heading direction. These body velocities were smoothed using a 15-frame gaussian-weighted moving average. An aggressively smoothed version of the body velocities was also generated using a 50-frame gaussian-weighted moving average instead; as described below, these velocities were used to identify turning bouts.

#### Data processing - Calcium imaging

Raw images in each time series were motion-corrected using the NoRMCorre software package^109^ to perform rigid subpixel registration of the jGCaMP7s channel. The CyRFP1 channel was shifted to match the jGCaMP7s channel. Regions of interest (ROIs) around the arbors of individual neurons were selected by thresholding on the ΔF/F signal computed over space in the time-series averaged image in the jGCaMP7s channel (i.e. for pixel i, F_i_/mean(F_all pixels_) - 1) and then by manually selecting the region of the thresholded mask to include. For each frame in the time series, intensity values for the pixels within each ROI were averaged and the mean background value (the average intensity in a region of the image without cells) was subtracted for each imaging frame within the time series. To correct for bleaching, the background-subtracted intensity time series for each ROI was then fit with the sum of two exponentials, and F(t)/F_0_ was calculated where F_0_ was the fitted value at each time t. F(t)/F_0_ was computed independently for the jGCaMP7s and CyRFP1 channels. To further account for motion artifacts, the jGCaMP7s signal was normalized to the CyRFP1 signal such that ΔF/F was (jGCaMP7s F(t)/F_0_) / (CyRFP1 F(t)/F_0_) - 1. The right-left difference and the right+left sum in ΔF/F was computed for the ROIs corresponding to the right and left cells for each trial.

#### Data processing - Tethered fly behavior

Raw FicTrac signals of the position of the ball in all three axes (yaw, pitch, and roll) were low-passed filtered at 40 Hz and then downsampled to 1000 Hz. These signals were then smoothed with a 200 ms time window Gaussian-weighted moving average. The fly’s velocity in each of the three axes (rotational, forward, and lateral) was computed by finding the derivative of the position signal. Rotational velocity was defined as the rate of change in the fly’s heading angle over time; this was taken as the opposite of the ball’s yaw velocity. Forward velocity was defined as the rate of change in position over time in the direction parallel to the fly’s anterior-posterior axis; this was taken as the opposite of the ball’s pitch velocity. Lateral velocity was defined as the rate of change in position over time in the direction orthogonal to the fly’s anterior-posterior axis; this was taken as the opposite of the ball’s roll velocity. The velocity signals were then smoothed with a 100 ms time window Gaussian-weighted moving average.

Using Animal Part Tracker (APT)^60^ running the Mixture Density Model (MDN) deep convolutional network, the tips of the 6 tarsi, the front of the head, the 3 midpoints between the 3 pairs of legs, and the tip of the abdomen, were tracked in each frame of the videos acquired from below the fly in our tethered fly behavioral setup. One network was trained for all of the tethered fly data, and it was a separate network from the one used for the freely walking flies. Tracked points were converted to positions in the fly’s coordinate frame and then to step parameters as described for the *Data analysis - Freely walking fly behavior* section, except for the following differences. The line used to define the anterior-posterior axis of the fly was fit directly to the head, the midpoints between each pair of legs, and the tip of the abdomen. The angle of the vector between the head and the abdomen points was not a readout of the fly’s heading as it was fixed; the FicTrac signal alone was used for heading. For each step, swing was defined as the half step with the shorter duration, and stance was the half step with the longer duration. Swing and stance were computed in this manner because it was not possible to determine whether the leg was moving or stationary relative to the ground when the ground (the ball) was also moving; however, the duration of the half-steps was a reliable readout of swing/stance as swing is always shorter than stance^65^.

For the calcium imaging experiments, time periods where the fly was not walking were determined by setting a threshold on the total speed across all three axes. This threshold was manually adjusted for each trial, and time points that fell below the threshold were preliminarily called not walking. The minimum duration of each walking or not-walking bout was 200 ms, and bouts that were too short were merged with their prior bout (i.e. the walk/not-walk call was flipped) until they exceeded the minimum duration.

For the optogenetic stimulation and electrophysiology experiments, time periods where the fly was not walking were determined by setting a threshold on the velocity of the two middle legs and on the total speed across all three body velocity axes. The thresholds were manually adjusted for each trial, and time points that fell below both thresholds were preliminarily called not walking. The minimum duration of each walking or not-walking bout was 200ms, and bouts that were too short were merged with their prior bout (i.e. the walk/not-walk call was flipped) until they exceeded the minimum duration.

#### Data processing - Electrophysiology

The time when each spike occurred was determined as the time of threshold crossing in the first derivative of the membrane potential. Threshold crossings that were less than 2 ms after the prior threshold crossing were excluded. The membrane potential threshold was determined manually for each trial. Spike rate was then computed as the inverse of the interspike interval, which was computed as the derivative of the vector of spike times.

#### Data processing - Connectomics

DNa02 was identified in the FAFB/FlyWire full *Drosophila* female brain connectome^23,27^ by manually comparing the morphology of every neuron that had its primary neurite in the appropriate tract with the light microscopy images of DNa02^28^. DNg13 was identified in FlyWire by first using natverse^110^ to register the light microscopy image of DNg13^28^ into the FAFB space to find the approximate location of the midline-crossing neurite and then by manually examining the morphology of every neuron that crossed the midline anywhere near that location. DNa02 was identified in FANC by the Jefferis lab. DNg13 was preliminarily identified in FANC by the Jefferis lab and that identification was then confirmed in this work by manual comparison of the morphology of every neuron that passed through the left half of the neck connective with the light microscopy images of DNg13^28^.

The connectivity of DNa02 and DNg13 in the brain was obtained from FlyWire^23,27,72,115,116^ using the BrainCircuits.io platform. Data was from the fruitfly_fafb_flywire_public API accessed on September 27, 2023. Each neuron presynaptic to DNa02 or DNg13 (minimum number of synapses = 10) was categorized into one of six categories by manual examination of its morphology. The categories were:

- Motor-associated - the input neuron arborizes predominantly in motor-associated brain regions (LAL, VES, IPS, SPS, EPA, or GNG)
- Descending neuron - the input neuron has a cell body in the brain and projects out of the brain through the neck connective
- Ascending neuron - the input neuron has a process in the neck connective and no cell body in the brain
- Visual - the input neuron has a dendritic arbor in a visual region (optic lobe, PVLP, AVLP, PLP, or AOTU)
- Superior brain - the input neuron’s dendrites are primarily in superior brain regions (SMP, SIP, SLP, MB, or LH)
- Central complex - the input neuron arborizes in the central complex (EB, PB, FB, NO, or AB; in practice, all of these neurons are PFL3).

Brain region abbreviations follow standard conventions^117^. Monosynaptic input neurons for the DNa02 and DNg13 cells in each hemisphere were considered to be shared if they made 10 or more synapses onto both neurons. They were considered to be unique inputs if they made 10 or more synapses onto one neuron but not the other.

The connectivity of DNa02 and DNg13 in the ventral nerve cord was obtained from FANC using the fancr package in natverse (https://github.com/flyconnectome/fancr). Many of the neurons downstream of DNa02 and DNg13 were proofread as part of a recent independent study^118^. The connectivity was last pulled on September 3, 2023. Motor neuron IDs were from the CAVEclient fanc_v4 tables: neck_motor_neuron_table_v0, haltere_motor_neuron_table_v0, wing_motor_neuron_table_v0, all_leg_motor_neuron_table_v0, and motor_neuron_table_v7^70^. Each neuron postsynaptic to DNa02 or DNg13 (minimum number of synapses = 10) was first categorized as likely leg or wing based on its morphology and basic connectivity:

- Leg - the neuron is a leg motor neuron, a leg premotor neuron, or arborizes in the leg neuromeres
- Wing - the neuron is a wing, haltere, or neck motor neuron or it arborizes in the dorsal neuropils and is not a leg premotor neuron

Neurons in the leg category were then divided into six further categories:

- Motor neuron - leg motor neuron
- Local premotor - interneuron whose arbors are restricted to one leg neuromere and that synapses onto at least one leg motor neuron
- Local - interneuron whose arbors are restricted to one leg neuromere and that does not synapse onto any leg motor neurons
- Intersegmental/bilateral premotor - neuron that arborizes in more than one leg neuromere, either bilaterally or intersegmentally, and synapses onto at least one leg motor neuron
- Intersegmental/bilateral - neuron that arborizes in more than one leg neuromere, either bilaterally or across segments, and that does not synapse onto any leg motor neurons
- Ascending neuron - neuron that has its cell body in the ventral nerve cord and sends a projection into the neck connective

Neurons in the wing category were then further divided into six categories:

- Wing motor neuron
- Haltere motor neuron
- Neck motor neuron
- Ascending neuron
- Unilateral interneuron - arborizes on one side of the dorsal neuropils
- Bilateral interneuron - arborizes on both sides of the dorsal neuropils

Monosynaptic output neurons for DNa02 and DNg13 were considered to be shared if they received 10 or more synapses from both DNs. They were considered to be unique outputs if they received 10 or more synapses from only one of the two neurons.

Neurotransmitter predictions were made by identifying the hemilineage each neuron belonged to based on anatomical similarity to the published descriptions of the neurons in each hemilineage and assigning the neuron’s neurotransmitter as the reported neurotransmitter for that hemilineage^71,103,104^.

#### Data analysis - Figures 1 and S1

Turning bouts were identified by finding peaks (using the MATLAB findpeaks function) in the extra smoothed version of the rotational velocity. The start and end of the bouts were defined as when the rotational velocity fell below 10°/s. Only bouts with a duration (end time – start time) of 0.3-0.7s were included in the subsequent analyses. Bouts where the fly turned left and bouts where the fly turned right were both extracted, with the left turn bouts being identified in the sign-inverted rotational velocity. Right and left turning bouts were combined by inverting the rotational velocity and step direction values and by swapping the left and right leg step parameter values for left turning bouts.

The step parameter values for each bout were the values during the step (for each leg) that occurred when the rotational velocity peaked. They were expressed as the change from the mean value across all flies when they were not turning (rotational speed < 10 °/s). In Figure 1C and Figure S1B, only bouts with a forward velocity of 10-15 mm/s at the peak in the rotational velocity were included. Bouts were then binned by their peak rotational velocity. In Figures 1E and S1C, only bouts with a forward velocity of 10-15 mm/s at the start of the bout and a peak rotational velocity of 50-150 °/s were included. Bouts were categorized as pivots if the difference between the forward velocity at the start of the bout and at the rotational velocity peak (peak – start) was less than −1 mm/s. Swerves were ones where the difference between the peak and the start was greater than 1 mm/s. Violin plots were generated using Violin Plot, from the MATLAB Central File Exchange (https://www.mathworks.com/matlabcentral/fileexchange/45134-violin-plot).

As step direction is a circular parameter, all descriptive and inferential statistics on it, in this and subsequent figures, were performed with the CircStat Toolbox^111^. For Figure S1C, Violin Plot was modified with CircStat functions to plot step direction appropriately.

For Figures 1C and S1B, two-way, unbalanced ANOVAs with leg identity and peak rotational velocity (including a no turning reference) as factors were performed and followed up with post-hoc Tukey-Kramer tests against reference for each rotational velocity and each leg.

For Figures 1E and S1C, except for step direction, two-way unbalanced ANOVAs with leg identity and pivot/swerve as factors were performed and followed up with post-hoc Tukey-Kramer tests comparing pivots and swerves for each leg. For step direction, parametric Watson-Williams multi-sample tests for equal means (CircStat Toolbox) were performed comparing pivot and swerve for each leg. Holm-Bonferroni correction for multiple comparisons was applied.

#### Data analysis - Figures 2 and S2

The heatmaps in Figure 2A were generated by plotting the mean right-left difference in ΔF/F for each of the rotational and forward velocity bins (30 bins each) for single example flies. ΔF/F was interpolated to 100 Hz. Only points from when the fly was walking were included in the heatmap. Data from transitions between walking and not walking periods were also excluded (0.14 s after the start of walking and 0.25 s before the end of walking). The time windows for exclusion were chosen to approximate the on kinetics of jGCaMP7s. Bins were colored gray if there were fewer than 20 data points.

For Figures 2B, S2A, and S2D, each fly’s mean rotational velocity (as the difference from that fly’s mean rotational velocity) was plotted against the right-left difference in ΔF/F (30 bins). For Figures S2B and S2E, each fly’s mean forward velocity (as the difference from that fly’s mean forward velocity) was plotted against the right+left sum in ΔF/F (30 bins). Similarly to Figure 2A, ΔF/F was interpolated to 100 Hz, and only points from when the fly was walking and not transitioning between walking and not walking were included. Bins with fewer than 50 data points were excluded. The mean across flies was also plotted. The Pearson correlation coefficient between the behavioral parameter and ΔF/F was computed for each fly and the mean ± SEM across flies is displayed in the figures.

For Figures 2C and 2D, linear filters relating the right-left difference in ΔF/F and the fly’s rotational velocity were estimated by computing the first-order Wiener kernel^119^ for each fly. Both ΔF/F and rotational velocity were interpolated to 100 Hz. In the frequency domain, the kernel F(ω) = I*(ω)R(ω)/I*(ω)I(ω), where I(ω) and R(ω) are the input and response, respectively. ω is the frequency and * indicates the complex conjugate. The kernels were computed in the frequency domain, low pass filtered according to c(ω) = e^(-abs(ω - f_cut_)/f_T_) for abs(ω) ≥ f_cut_. f_cut_ was 4Hz and f_T_ was 1Hz. For Figure 2C, ΔF/F was the input and rotational velocity was the response. For Figure 2D, rotational velocity was the input and ΔF/F was the response. The kernels were computed for each fly using all data, regardless of whether the fly was walking or not, and then averaged across flies. The time axis in Figure 2C was inverted so negative time values were when ΔF/F preceded behavior for both types of filters.

For Figures S2C and S2F, the mean right+left sum in ΔF/F was calculated for each fly when it was walking and when it was not walking. Paired t-tests for a difference between walking and not walking (paired by fly) were performed for each cell type.

#### Data analysis - Figures 3 and S3

Figure 3A displays the number of neurons that are presynaptic to DNa02 alone, DNg13 alone, or shared between the two for each hemisphere. Figure S3B shows the total number of synapses made onto the DNs by the neurons in each of those categories.

In Figure 3B, monosynaptic inputs of DNa02 and DNg13 were grouped by category; each neuron was treated equally and the plot expresses category membership as the percent of the total number of input neurons.

Figure S3C totals up the number of synapses made onto the DNs by neurons in each category, and the plot displays the number of synapses each category contributes, as a percent of the total number of synapses.

For DNa02 in Figure 3D, the duration of optogenetic stimulation was 200 ms, and the illumination power was 0.09 mW/mm^2^. For DNg13, the duration of the current injection was 1 s, and the amplitude was 100 pA.

#### Data analysis - Figures 4 and S4

For Figures 4A-C and S4, the duration of optogenetic stimulation was 200 ms and the illumination power was 0.09 mW/mm^2^. For Figures 4D-F and S4D, the duration of current injection was either 0.5 or 1 s. For hyperpolarization (−75 pA current injection step), only trials where the spike rate was suppressed below 5 spikes/s were analyzed; for depolarization (100 pA current injection step), only trials where the spike rate was increased to above 130 spikes/s were analyzed. For all panels of Figures 4 and S4, stimulation trials were only included if the fly maintained an average forward velocity of 1 mm/s or greater from 200 ms before the start of the stimulation to 200 ms after the end of stimulation. Matched non-stimulated epochs for each fly were defined as a time window that matched the stimulation duration in the middle of each inter-stimulation period and that met the forward velocity criteria.

For Figures 4A, 4D, and S4A, the plotted mean rotational velocity for each fly was the mean cumulative change in heading direction across perturbation trials minus the mean value during non-stimulated epochs in the same fly, divided by the time window of consideration (500 ms from the start of stimulation). The plotted mean forward velocity for each fly was computed identically, except with the mean cumulative forward displacement instead of heading direction. For Figures 4A and S4A, two-sample t-tests were performed comparing ±CsCh for the difference in rotational velocity and the difference in forward velocity. For Figure 4D, two-sample t-tests comparing hyperpolarization or depolarization trials with no-stimulation trials were performed.

In Figures 4B, 4C, 4E, and 4F, the change in the step parameter was measured as the mean value during perturbation, minus the mean value during non-stimulated epochs in the same fly. The change was measured using steps that fell between the start of the stimulation to 0.1 s after the end of stimulation. These time windows are different from those of Figure 4A and 4D because step parameters are computed only once per step cycle.

For Figures 4B, 4C, S4B, and S4C, unbalanced 2-way ANOVAs with CsCh expression and leg identity as factors were performed and followed up with post-hoc Tukey-Kramer tests comparing ±CsCh for each leg.

In Figure 4E, unbalanced 3-way ANOVA with depolarization/hyperpolarization, leg identity, and fly identity as factors was performed. The effect of hyperpolarizing DNg13 versus depolarizing DNg13 was compared using post-hoc paired t-tests. For Figure 4F, unbalanced 3-way ANOVA with depolarization/hyperpolarization, leg identity, and fly identity as factors, without interaction, using contralateral legs only was performed.

In Figure S4D, parametric two-way ANOVAs for circular data with interaction were performed, with leg identity and ±CsCh or depolarization/hyperpolarization as factors. For experiments with significant differences (left and middle), post-hoc parametric Watson-Williams two-sample tests for equal means, comparing ±CsCh or depolarization/hyperpolarization conditions for each leg were performed. Holm-Bonferroni correction for multiple comparisons was performed.

#### Data analysis - Figures 5 and S5

Behavioral variables were compared with normalized DN spike rate 100 ms prior. Normalized spike rate was computed by subtracting the mean spike rate when the fly was not walking (which is typically very low in these DNs) and then dividing by the 95^th^ percentile value of the resulting distribution. For the stride parameters, continuous estimates were used rather than their values per step. Ipsilateral/contralateral stride length was defined as the average of the mean subtracted stride length for each of the three legs ipsilateral/contralateral to the recorded DN soma. Ipsilateral/contralateral return stroke position was the average of the return stroke position for all ipsilateral/contralateral legs, with values from each leg centered by subtracting the mean for that leg. Ipsilateral/contralateral power stroke position was the average of the power stroke position for all ipsilateral/contralateral legs, with values from each leg centered by subtracting the mean for that leg.

For Figures 5B-D, S5B, and S5C, the mean in the behavioral variable was computed for each fly across 50 bins of normalized DN spike rate. Only data from times when the mid and hind legs were walking forwards (based on step direction) were included. Data were interpolated to 250 Hz. Bins with fewer than 50 data points were excluded from the mean. The Pearson correlation coefficient between the behavioral parameter and the spike rate was computed for each fly and the mean ± SEM across flies is displayed in the figure panels. For Figures 5E-G, the data was further split into times when the fly’s rotational speed was less than 25 °/s (not turning) and times when it was greater than or equal to 25 °/s (turning). The Pearson correlation coefficient was computed between the behavioral variable and the spike rate for these two conditions. Paired t-tests were performed.

For Figure S5A, the mean spike rate was computed for each fly when it was not walking and when it was walking with a rotational velocity of less than 0 °/s ipsiversive to the recorded DN soma (i.e. the fly was not turning ipsilaterally). Paired t-tests were performed.

For Figure S5D, the Pearson correlation coefficient was computed between each pair of behavioral variables that was plotted in Figures 5B-D, S5B, and S5C. Only data from times when the mid and hind legs were walking forwards were included. The correlation was computed for each fly, and the mean across flies was reported.

#### Data analysis - Figure 6

For Figure 6A, the linear filters relating rotational velocity or stride length (ipsilateral for DNa02 and contralateral for DNg13) with DN spike rate were computed by first computing the cross-correlation between the behavioral variable and the spike rate as well as the autocorrelation of the spike rate, transforming them into the frequency domain, dividing them, low pass filtering them, and then transforming them back into the time domain. This effectively computes the first-order Wiener kernel. Only data from times when the mid and hind legs were walking forwards were included. The cross-correlation and the autocorrelation were computed by calculating the Pearson correlation coefficient at every time lag between −1 and 1s, in 1 ms increments. The data was low pass filtered according to c(ω) = e^(-abs(ω - f_cut_)/f_T_) for abs(ω) ≥ f_cut_. f_cut_ was 2 Hz and f_T_ was 0.5 Hz. Cells for which the cross-correlation did not have a defined peak were excluded, as they did not produce meaningful filters. The filter relating the ipsilateral stride length and DNa02 spike rate was sign inverted to make comparison with the DNg13 filter easier. The peak time was the time at which the maximum in the filter occurred. One-sample t-tests were performed to determine whether the peak times were significantly different from 0.

Figure 6B plots the autocorrelation for DNa02 and DNg13 spike rate. Only data from times when the mid and hind legs were walking forwards were included. The full-width at half-maximum of the autocorrelation function for each fly was determined by finding the time window when the autocorrelation was greater than or equal to 0.5. A two-sample t-test was performed to determine whether the difference in the full-width at half-maximum between DNa02 and DNg13 was significant.

For Figure 6C, turning bouts were identified in the freely walking fly data as described for Figure 1. Bouts were included if they had an initial forward velocity of 10-15 mm/s, a peak rotational velocity of 50-150 °/s, and a duration of 0.3-0.7s. The continuous estimate of stride length from 0.5 s before the peak in yaw velocity to 0.5 s after the peak in yaw velocity was extracted for each leg in each bout; this was interpolated to 100 Hz. This was then normalized by subtracting the minimum stride length during the snippet and dividing by the range (max - min). The mean across bouts was then computed for each leg. For plotting, the mean was again normalized to 0-1 by subtracting the minimum and dividing by the range. The full-width at half-maximum was computed on the normalized stride length for each bout. The maximum was found in the stride length within ±0.2 s of the peak in rotational velocity, and the duration that the stride length spent greater than the half-max value was determined.

In Figure 6D, the phase of the stride cycle was computed as the average phase across all six legs, with the phases of all the legs aligned by subtracting the mean offset relative to a reference leg (ipsilateral middle leg for DNa02 and contralateral middle leg for DNg13). Each spike was assigned to a phase bin (72 bins) based on the phase 100 ms prior. Spike counts per bin were normalized by the time spent at each phase and then expressed as the difference from the mean across all phases. All spikes when the fly was walking were used.

#### Data analysis - Figures 7 and S6

In Figures 7A and 7B, monosynaptic outputs of DNa02 and DNg13 were grouped by category; like for Figure 3B, each neuron was treated equally, and the plot expresses category membership as the percent of the total number of output neurons in the leg neuromeres. In Figures S6E and S6F, the plots on the right are similar, except that Figure S6E categorizes all output neurons, and Figure S6F looks at outputs in the dorsal neuropil regions, specifically. Figures S6B and S6C and the left plots in Figures S6E and S6F sum the number of synapses made by the neurons in each category and express them as a percent of the total number of synapses (like Figure S3C).

Figure 7C displays the number of neurons that are postsynaptic to DNa02 alone, DNg13 alone, or shared between the two for each hemisphere. Figure S6D shows the total number of synapses made onto neurons in each of those categories.

Figure 7D shows the monosynaptic and disynaptic connections between DNa02/DNg13 and the T1 leg motor neurons (minimum number of synapses = 15). The graphs for DNa02 and DNg13 are plotted separately, so there are motor neurons that are repeated between the two. Connections between premotor neurons were not displayed. Leg motor neurons were labeled based on their inferred effect on the leg^70^. Neurotransmitter labels reflect machine vision predictions^72^ (for DNs) or hemilineage membership^71,103,104^ (for premotor cells). Neurotransmitters were deemed excitatory or inhibitory based on the effect of their ionotropic receptors; glutamate was deemed inhibitory as the inhibitory glutamate-gated chloride channel GluClɑ^120,121^ is widely expressed in the ventral nerve cord^122^.

Figures 7E and 7F show the premotor and motor neurons downstream of DNg13 that were hypothesized to be active during the power (Figure 7E) and return (Figure 7F) phases of the stride cycle. Neurons were hypothesized to be active if their inferred effect on the leg (accounting for the sign of the connections) was consistent with the results of DNg13 perturbation (Figures 4 and S4) during that phase.

### Quantification and statistical analysis

All statistical methods used in the paper are described briefly in the figure legends and in detail in the method details. The p-values for all statistical tests are in Table S1. Statistical significance was defined at ɑ = 0.05. When appropriate, Holm-Bonferroni correction was applied for multiple comparisons. Definitions of sample size, measures of center and dispersion, and precision measures are also indicated in figure legends and method details. Statistics were computed using MATLAB.

**Figure S1:**
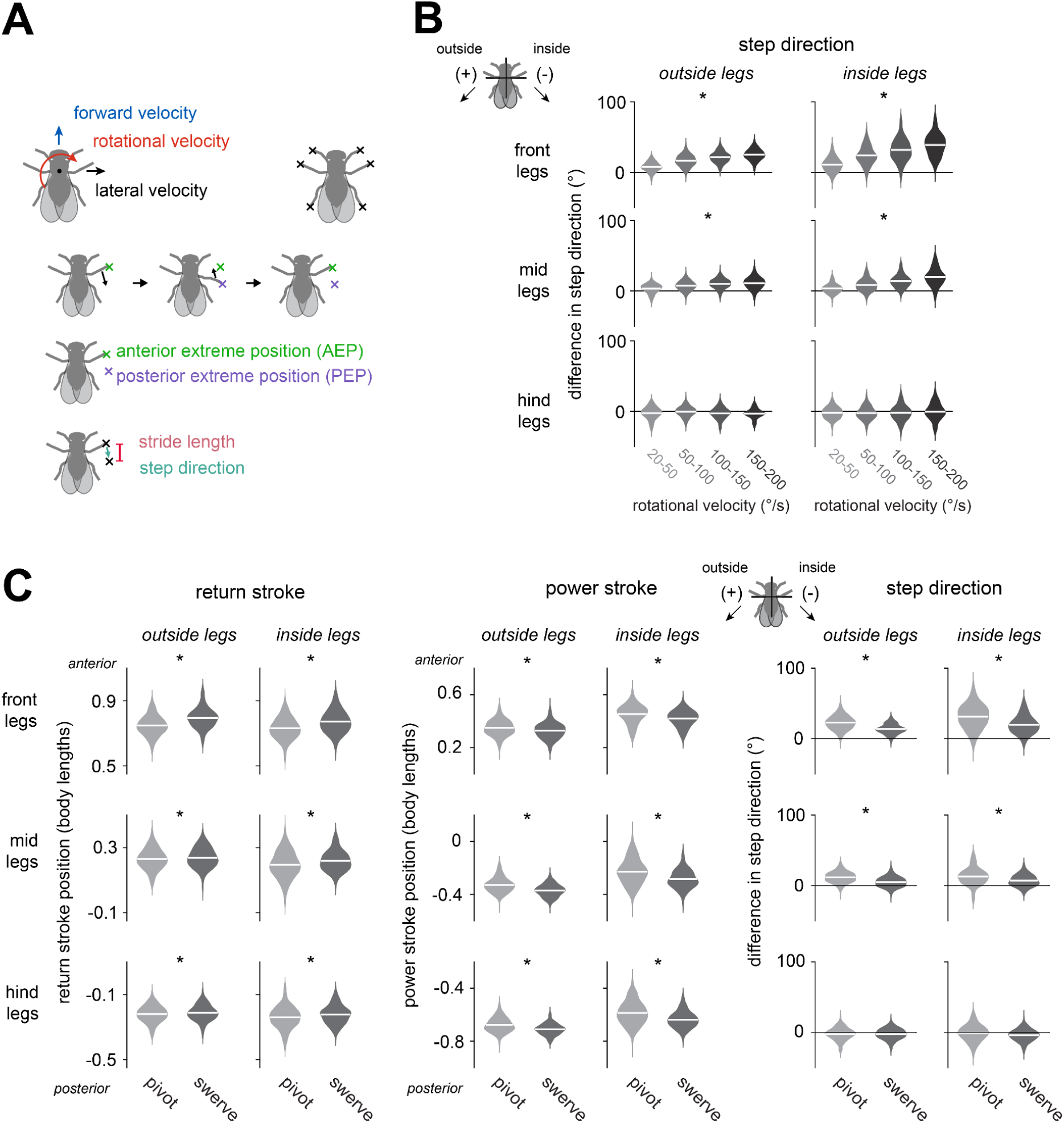
Modulations of step parameters during steering in freely walking flies. (A) *Top*: schematic of the walking features tracked: the velocity in three body axes and the positions of the tips of the 6 legs. *Bottom:* during a step cycle, the tip of the leg moves posterior relative to the body and then anterior, defining the anterior and posterior extreme positions (AEP and PEP). The stride length is the anterior-posterior distance between these two positions and the step direction is the angle of the vector from the AEP to the PEP. (B) When flies steer, they modulate the step direction of the front and mid legs (parametric Watson-Williams multi-sample test for equal means: out-front: p=0.0277; out-mid: p=1.94×10^−6^; out-hind: p=0.943; in-front: p=5.55×10^−16^; in-mid: p=1.42×10^−4^; in-hind: p=0.963). * marks legs with significant differences. For each turning bout, we measured the bout’s peak rotational velocity, as well as the step direction for the step that occurred at that peak. Step directions are expressed as the change from the mean when the fly was not turning. n = 138 (55), 454 (68), 434 (68), and 151 (50) bouts (flies) for 20-50, 50-100, 100-150, and 150-200 °/s, respectively. (C) Pivots and swerves differentially modulate the return stroke position, the power stroke position, and the step direction. The return and power stroke positions are plotted as positions in the anterior-posterior axis, and step direction is plotted as the difference from the mean when the fly was not turning (two-way ANOVA with legs and pivot/swerve as factors: return stroke: pivot/swerve p=3.68×10^−15^, interaction between pivot/swerve and leg identity: p=0.0198; power stroke: pivot/swerve p=1.75×10^−27^, interaction between pivot/swerve and leg identity: p=0.794; post-hoc Tukey-Kramer tests show a significant difference between pivots and swerves (Table S1); step direction: parametric Watson-Williams multi-sample test for equal means, comparing pivot/swerve for each leg, out-front: p = 2.22×10^−16^, out-mid: p=2.22×10^−16^; out-hind: p=0.445; in-front: p=4.13×10^−12^; in-mid: p=9.94×10^−5^; in-hind: p = 0.140). n = 389 (64) and 155 (53) bouts (flies) for pivots and swerves, respectively.

**Figure S2:**
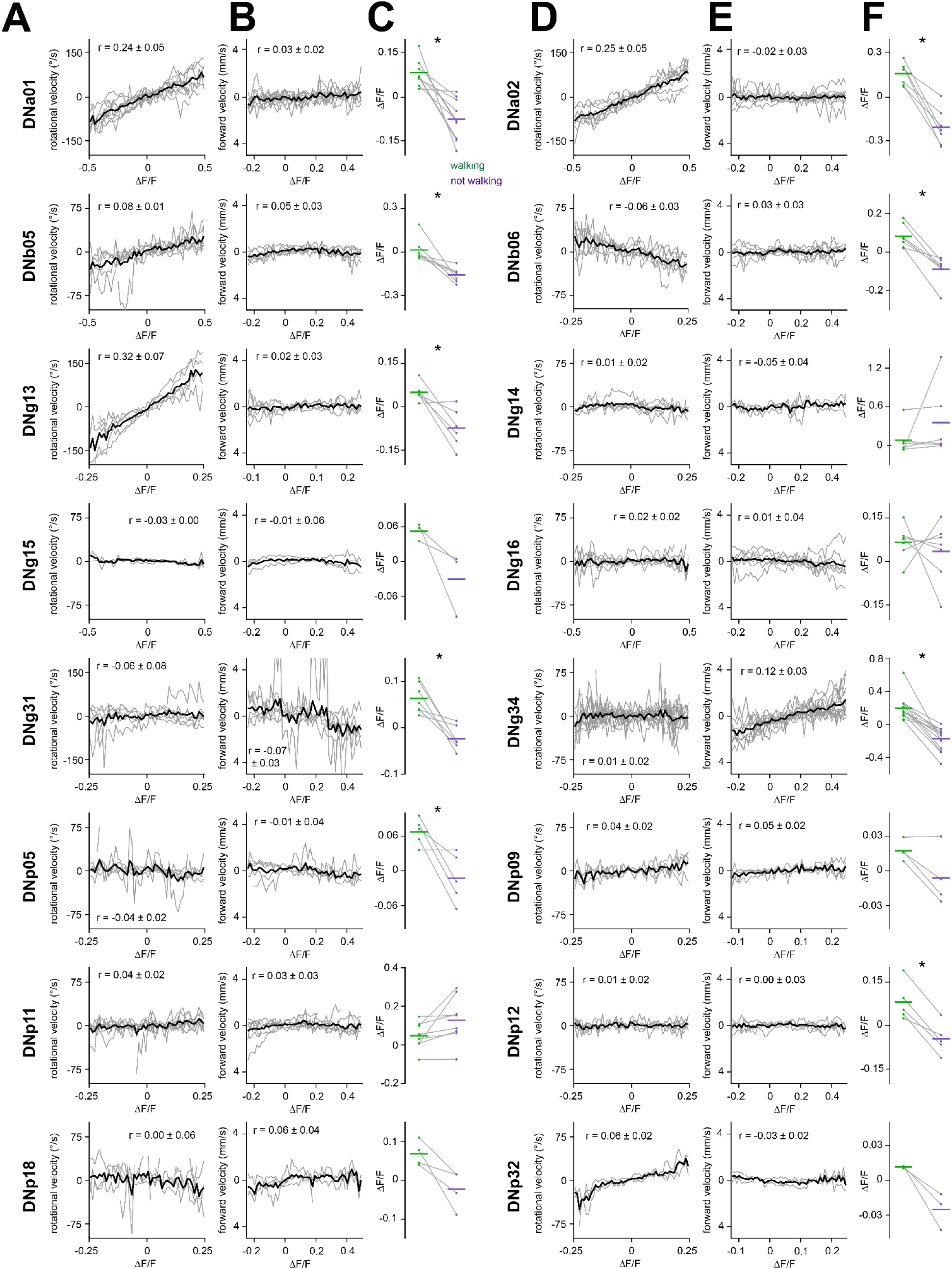
Correlations between the activity of all DN types imaged and walking. Each DN type imaged is represented by a single right-left pair of neurons. The three plots for each cell type are: *(A and D)* rotational velocity vs. the right cell – left cell difference in ΔF/F. Individual flies are in gray, and the mean across flies is in black. *(B and E)* forward velocity vs. the right cell + left cell sum in ΔF/F. *(C and F)* the mean right + left sum in ΔF/F during time periods when the fly is walking or not walking. The mean activity is higher when the fly was walking than when it was not walking for many DN types (* marks significant differences, see Table S1). Dots are individual flies, and the horizontal line is the mean across flies. Gray lines connect points from the same fly. n=10 flies for DNa01; n=8 flies for DNa02; n=7 flies for DNb05; n=7 flies for DNb06; n=5 flies for DNg13; n=6 flies for DNg14; n=3 flies for DNg15; n=7 flies for DNg16; n=7 flies for DNg31; n=14 flies for DNg34; n=5 flies for DNp05; n=4 flies for DNp09; n=5 flies for DNp11; n=5 flies for DNp12; n=5 flies for DNp18; n=4 flies for DNp32.

**Figure S3:**
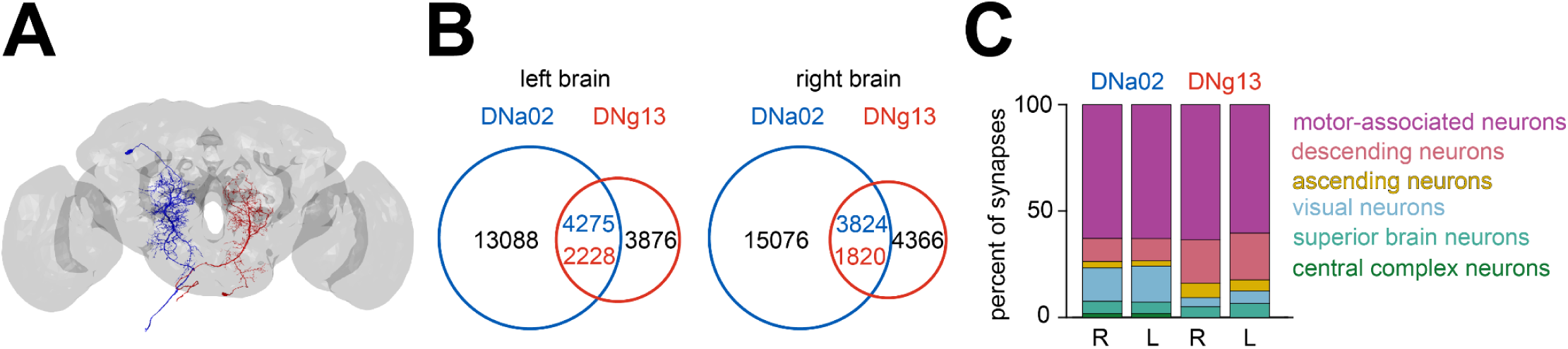
Steering descending neurons have distinct inputs. (A) Image shows the morphology of the left DNa02 and the right DNg13 in the brain, from the full female brain connectome^23,27^. (B) The number of synapses made onto the DNs by their shared or unique presynaptic cells. For shared inputs, the number of synapses is colored based on the postsynaptic DN. Shared presynaptic cells contribute a smaller fraction of the DN’s inputs than their unique presynaptic partners. (C) Stacked bar charts summarize the number of synapses made by the cells in each category onto each DN in the brain, expressed as a percentage of all presynapses.

**Figure S4:**
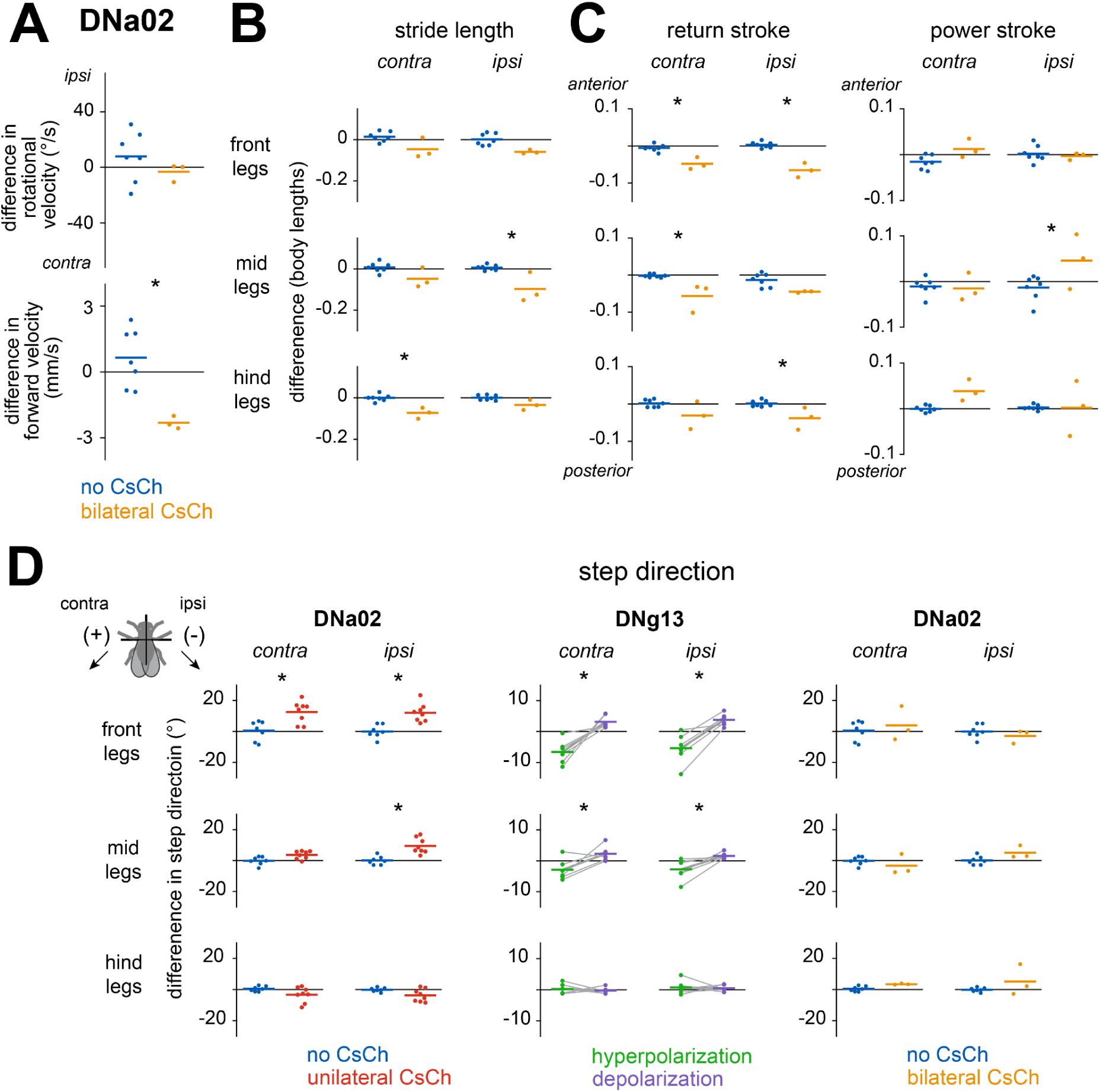
Perturbing steering-related descending neurons. (A) Difference in rotational and forward velocity during bilateral optogenetic activation of DNa02 with CsChrimson (CsCh). Dots are individual flies, and lines are the means across flies. Ipsiversive (positive) and contraversive (negative) were randomly assigned for each fly. The effect of optogenetic stimulation is significant for forward velocity (p=0.00564) but not for rotational velocity (p=0.341; two-sample t-tests, n=3 flies (CsCh+) and 7 flies (no CsCh)). * marks significant changes upon stimulation. (B) Bilateral optogenetic stimulation changes the stride length across legs (two-way ANOVA with CsCh expression and leg identity as factors, ±CsCh: p=1.25×10^−10^; interaction between CsCh expression and leg identity: p=0.253 (* marks legs with significant differences in post-hoc tests, see Table S1)). (C) Bilateral optogenetic stimulation of DNa02 shortens the return stroke but has little consistent effect on the power stroke (two-way ANOVA with CsCh expression and leg identity as factors: return stroke: ±CsCh: p=5.37×10^−13^; interaction between CsCh expression and leg identity: p=0.180; power stroke: ±CsCh: p=0.00714; interaction: p=0.0414 (see Table S1 for post-hoc tests)). (D) *Left:* unilateral activation of DNa02 with optogenetic stimulation of CsCh produces changes in the step direction of the front legs and the ipsilateral mid leg (parametric two-way ANOVA for circular data with CsCh expression and leg identity as factors, ±CsCh: p=5.37×10^−13^; interaction between CsCh expression and leg identity: p=0.180 (see Table S1 for post-hoc tests)). *Middle:* unilateral hyperpolarization and depolarization of DNg13 produce changes in the step direction of the front and mid legs (parametric two-way ANOVA for circular data with depolarization/hyperpolarization and leg identity as factors: depolarization/hyperpolarization: p=9.82×10^−10^; interaction between depolarization/hyperpolarization and leg identity: p=8.14×10^−6^ (see Table S1 for post-hoc tests)). *Right:* bilateral activation of DNa02 does not change the step direction of any of the legs (parametric two-way ANOVA for circular data with CsCh expression and leg identity as factors, ±CsCh: p=0.176; interaction between CsCh expression and leg identity: p=0.232). Changes in step direction are expressed as the difference from the step direction during matched periods without stimulation. Dots are individual flies, and the horizontal lines are the means across flies. Gray lines connect points from the same fly.

**Figure S5:**
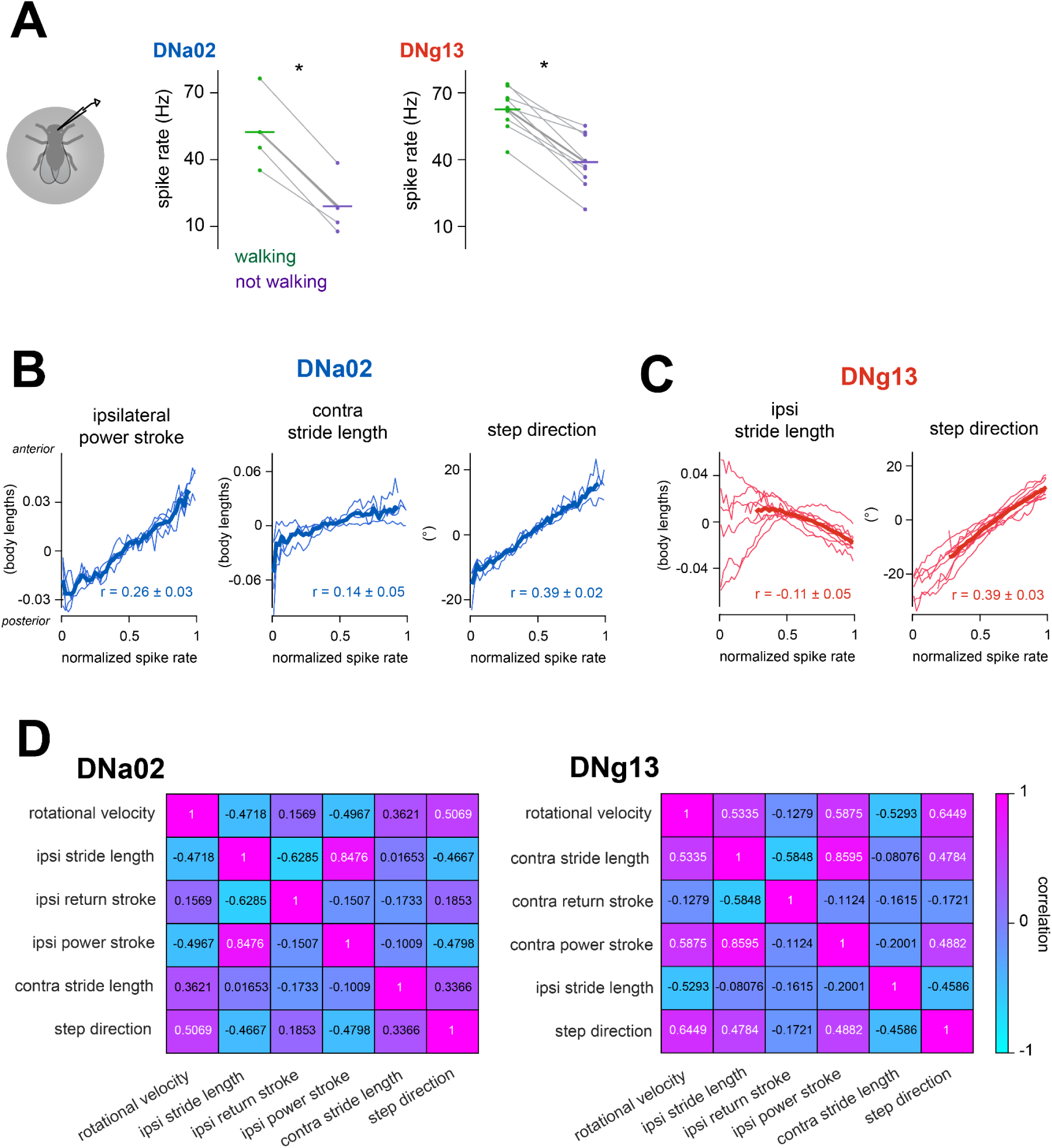
Comparing descending neuron activity with behavior. (A) DNa02 and DNg13 spike rates are greater when the fly is walking than when the fly is not walking. Plot shows the mean DNa02/DNg13 spike rate when the fly was walking and not making ipsiversive turns (rotational velocity < 0 °/s) or when the fly was not walking. Dots are individual flies, and horizontal lines are the means across flies. Points from the same fly are connected with gray lines (paired t-test on spike rate, walking vs. not walking, DNa02: p=0.00229; DNg13: p=4.59×10^−5^). (B) Ipsilateral power stroke position, contralateral stride length, and front-leg step direction are related to DNa02 spike rate. Thin lines are mean values for individual flies; thick lines are averages across flies (n=4 cells in 4 flies). The mean ± SEM across flies of the correlation between the behavioral parameters and the spike rate is also shown. (C) Same but for DNg13 spike rate and ipsilateral stride length and front-leg step direction (n=9 cells in 9 flies). (D) Many behavioral parameters are correlated during turning. Mean pairwise correlations (Pearson correlation coefficient) between behavioral parameters that were correlated with DNa02 and DNg13 spike rate, in the same flies that the spike rate-behavior correlations were computed.

**Figure S6:**
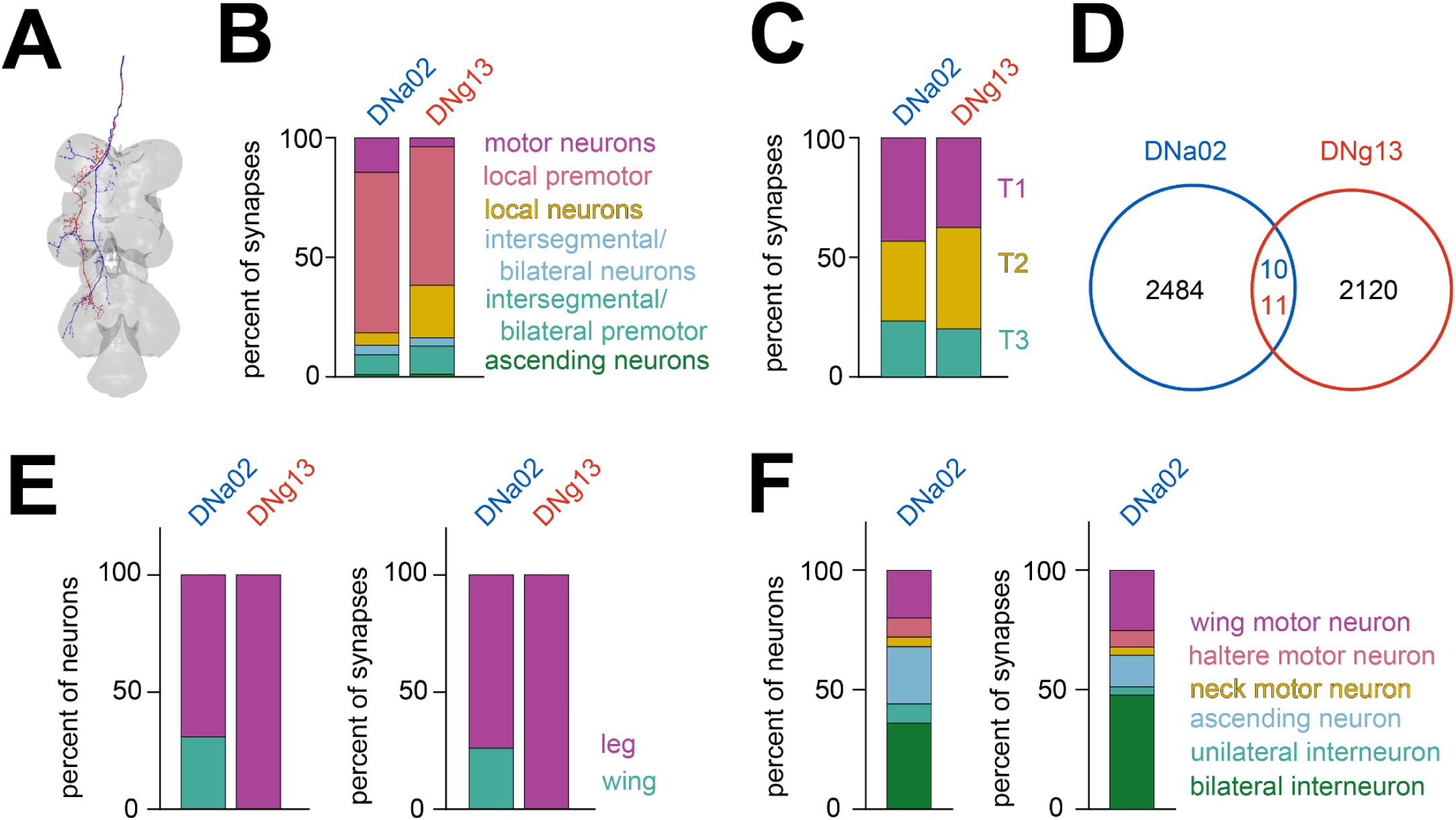
Connectivity of steering descending neurons in the ventral nerve cord. (A) Image shows the morphology of the DNa02 axon and the DNg13 axon on the left side of the ventral nerve cord^25^. These axons come from the DNa02 in the left hemisphere of the brain and the DNg13 in the right hemisphere of the brain. (B) Number of output synapses from DNa02 and DNg13 onto cells in each category in the leg neuromeres of the ventral nerve cord, expressed as a percentage of all postsynapses made in the leg neuromeres. Note that DNa02 synapses onto neurons in the dorsal neuropils (associated with the wings, halteres, and neck), and those are excluded from this plot. (C) The number of synapses made onto neurons in each of the three leg neuromeres (T1, T2, T3), expressed as a percentage of all postsynapses made in the leg neuromeres. Intersegmental neurons are assigned to a segment based on their cell body locations. (D) The number of synapses made onto shared and unique postsynaptic targets of each DN. For shared targets, the number of synapses made is colored based on the presynaptic DN. DNa02 and DNg13 axons projecting to the left side of the ventral nerve cord make very few synapses onto the same cells. (E) *Left:* The percent of all monosynaptic output neurons in the ventral nerve cord that likely belong to leg or wing circuitry. *Right:* the number of synapses made onto leg or wing circuitry neurons, expressed as a percentage of the total synapses made by these DNs. (F) Number of neurons (*left*) and the number of synapses made onto these neurons (*right*) for the monosynaptic output neurons classified as belonging to the wing circuitry.

**Table S1:** Statistics.

| Figure | Test | p-values |
| --- | --- | --- |
| Figure 1C | Two-way ANOVA with legs and rotational velocity as factors | rotational velocity: $p = 4.87 \times 10^{-77}$; leg identity: $p = 0$; interaction between leg identity and rotational velocity: $p = 0$ |
| Figure 1C | Post-hoc Tukey-Kramer tests against reference for each rotational velocity (20-50°/s, 50-100°/s, 100-150°/s, 150-200°/s) | In ascending order of rotational velocity: out-front [0.0734, $6.67 \times 10^{-12}$, 0, $1.47 \times 10^{-8}$]; out-mid [0.0409, $7.15 \times 10^{-18}$, 0, 0]; out-hind [0.807, $1.68 \times 10^{-4}$, 0.00437, $7.18 \times 10^{-10}$]; in-front [0.00167, 0, 0, 0]; in-mid [1.00, 0, 0, 0]; in-hind [0.627, 0, 0, 0] |
| Figure 1E | Two-way ANOVA with legs and pivot/swerve as factors | pivot/swerve: $p = 6.11 \times 10^{-31}$; leg identity: $p = 4.96 \times 10^{-66}$; interaction: $p = 0.738$ |
| Figure 1E | Post-hoc Tukey-Kramer tests comparing pivot and swerve for each leg | out-front: $p = 4.28 \times 10^{-8}$; out-mid: $p = 0.000744$, out-hind: $p = 0.000200$; in-front: $p = 0.000243$; in-mid: $p = 5.29 \times 10^{-5}$; in-hind: $p = 0.00524$ |
| Figure 4A | Two-sample t-tests comparing CsCh+ and no-CsCh | rotational velocity: $p = 0.00141$; forward velocity: $p = 0.802$ |
| Figure 4B | Two-way ANOVA with CsCh expression and leg identity as factors | ±CsCh: $p = 2.85 \times 10^{-4}$; leg identity: $1.98 \times 10^{-6}$; interaction between leg identity and ±CsCh $p = 8.78 \times 10^{-6}$ |
| Figure 4B | Post-hoc Tukey-Kramer tests comparing CsCh+ and no-CsCh for each leg | c-front: $p = 1.00$; c-mid: $p = 0.937$; c-hind: $p = 0.996$; i-front: $p = 0.0408$; i-mid: $p = 0.000181$; i-hind: $p = 0.0506$. |
| Figure 4C (left) | Two-way ANOVA with CsCh expression and leg identity as factors | ±CsCh: $p = 4.23 \times 10^{-6}$; leg identity: $p = 1.27 \times 10^{-8}$; interaction between leg identity and ±CsCh: $p = 2.46 \times 10^{-8}$ |
| Figure 4C (left) | Post-hoc Tukey-Kramer tests comparing CsCh+ and no-CsCh for each leg | c-front: $p = 0.987$; c-mid: $p = 0.957$; c-hind: $p = 1.00$, i-front: $p = 5.08 \times 10^{-7}$; i-mid: $p = 0.00716$; i-hind: $p = 0.00118$ |
| Figure 4C (right) | Two-way ANOVA with CsCh expression and leg identity as factors | ±CsCh: $p = 0.807$; leg identity: $p = 0.0779$; interaction between leg identity and ±CsCh: $p = 0.0354$ |
| Figure 4D | Two-sample t-tests comparing depolarization or hyperpolarization with no-stimulation trials | rotational velocity, hyperpol: $p = 0.00257$; rotational velocity, depol: $p = 0.0404$; forward velocity, hyperpol: $p = 0.828$; depol: $p = 0.695$ |
| Figure 4E | 3-way ANOVA with depol/hyperpol, leg identity, and fly identity as factors | fly identity: $p = 0.0661$; leg identity: $p = 0.262$; depol/hyperpol: $p = 0.000149$; fly identity and leg identity interaction: $p = 0.513$; fly identity and depol/hyperpol interaction: $p = 0.502$; leg identity and depol/hyperpol interaction: $p = 0.000501$ |
| Figure 4E | Post-hoc, paired t-tests comparing depol and hyperpol for each leg, with Holm-Bonferroni correction for multiple comparisons | c-front: $p = 0.0340$; c-mid: $p = 0.00136$ *; c-hind: $p = 0.00200$ *; i-front: $p = 0.127$; i-mid: $p = 0.786$; i-hind: $p = 0.449$ (where * denotes significance after multiple comparisons correction, $\alpha = 0.05$) |
| Figure 4F (left) | 3-way ANOVA depol/hyperpol, leg identity, and fly identity as factors, without interaction, using contralateral legs only | depol/hyperpol: $p = 7.84 \times 10^{-4}$; leg identity: $p = 0.696$; fly identity: $p = 0.292$ |
| Figure 4F (right) | 3-way ANOVA depol/hyperpol, leg identity, and fly identity as factors, | depol/hyperpol: $p = 5.28 \times 10^{-4}$; leg identity: $p = 0.252$; fly identity: $p = 0.0175$ |
|  | without interaction, using contralateral legs only |  |
| Figure 5E (right) | paired t-test on difference in correlation coefficients, turning versus not-turning | $p = 0.0109$ |
| Figure 5F (right) | paired t-test on difference in correlation coefficients, turning versus not-turning | $p = 0.00836$ |
| Figure 5G (right) | paired t-test on difference in correlation coefficients, turning versus not-turning | $p = 0.00767$ |
| Figure 6A (left) | one-sample t-test, time of peak different from 0 | DNa02: $p = 2.85 \times 10^{-4}$; DNg13: $p = 5.36 \times 10^{-4}$ |
| Figure 6A (right) | one-sample t-test, time of peak different from 0 | DNa02: $p = 0.00250$; DNg13: $p = 7.24 \times 10^{-4}$ |
| Figure 6B | two-sample t-test on the full-width at half-maximum for DNa02 vs. DNg13 | $p = 0.00520$ |
| Figure 6C (right) | paired t-test for each pair of legs comparing the ipsilateral and contralateral full-width at half-maximum | front: $p = 1.31 \times 10^{-76}$; mid: $p = 8.08 \times 10^{-84}$; hind: $p = 4.95 \times 10^{-93}$ |
| Figure S1B | For each leg, parametric Watson-Williams multi-sample test for equal means | out-front: $p = 0.0277$; out-mid: $p = 1.94 \times 10^{-6}$; out-hind: $p = 0.943$; in-front: $p = 5.55 \times 10^{-16}$; in-mid: $p = 1.42 \times 10^{-4}$; in-hind: $p = 0.963$ |
| Figure S1C (left) | Two-way ANOVA with legs and pivot/swerve as factors | pivot/swerve: $p = 3.68 \times 10^{-15}$, leg identity: $p = 1.01 \times 10^{-17}$, interaction: $p = 0.0198$ |
| Figure S1C (left) | Post-hoc Tukey-Kramer tests comparing pivots and swerves for each leg | out-front: $p = 4.28 \times 10^{-8}$; out-mid: $p = 0.000744$; out-hind: $p = 0.000200$; in-front: $p = 0.000243$; in-mid: $p = 5.29 \times 10^{-5}$; in-hind: $p = 0.00524$ |
| Figure S1C (middle) | Two-way ANOVA with legs and pivot/swerve as factors | pivot/swerve $p = 1.75 \times 10^{-27}$, leg identity $p = 0$, interaction: $p = 0.794$ |
| Figure S1C (middle) | Post-hoc Tukey-Kramer tests comparing pivots and swerves for each leg | out-front: $p = 0.00220$; out-mid: $p = 4.75 \times 10^{-6}$; out-hind: $p = 0.00120$; in-front: $p = 0.0235$; in-mid: $p = 1.54 \times 10^{-5}$; in-hind: $p = 0.00143$ |
| Figure S1C (right) | For each leg, parametric Watson-Williams multi-sample test for equal means | out-front: $p = 2.22 \times 10^{-16}$, out-mid: $p = 2.22 \times 10^{-16}$; out-hind: $p = 0.445$; in-front: $p = 4.13 \times 10^{-12}$; in-mid: $p = 9.94 \times 10^{-5}$; in-hind: $p = 0.140$; |
| Figures S2C and S2F | paired t-tests comparing walking and not-walking | DNa01: $p = 0.000595$; DNa02: $p = 0.000558$; DNb05: $p = 0.00644$; DNb06: $p = 0.00301$; DNg13: $p = 0.0138$; DNg14: $p = 0.302$; DNg15: $p = 0.167$; DNg16: $p = 0.607$; DNg31: $p = 0.00370$; DNg34: $p = 5.60 \times 10^{-6}$; DNp05: $p = 0.0290$; DNp09: $p = 0.0715$; DNp11: $p = 0.0520$; DNp12: $p = 0.00234$; DNp18: $p = 0.0503$; DNp32: $p = 0.0628$ |
| Figure S4A | Two-sample t-tests comparing CsCh+ and no-CsCh | rotational velocity: $p = 0.341$; forward velocity: $p = 0.00564$ |
| Figure S4B | Two-way ANOVA with CsCh expression and leg identity as factors | ±CsCh: $p = 1.25 \times 10^{-10}$; leg identity: $p = 0.201$; interaction between leg identity and ±CsCh: $p = 0.253$ |
| Figure S4B | Post-hoc Tukey-Kramer tests comparing CsCh+ and no-CsCh for each leg | c-front: $p = 0.112$; c-mid: $p = 0.197$; c-hind: $p = 0.0219$,<br>i-front: $p = 0.0998$; i-mid: $p = 1.44 \times 10^{-4}$; i-hind: $p = 0.800$ |
| Figure S4C (left) | Two-way ANOVA with CsCh expression and leg identity as factors | ±CsCh: $p = 5.37 \times 10^{-13}$; leg identity: $p = 0.219$; interaction between leg identity and ±CsCh: $p = 0.180$ |
| Figure S4C (left) | Post-hoc Tukey-Kramer tests comparing CsCh+ and no-CsCh for each leg | c-front: $p = 0.0174$; c-mid: $p = 7.49 \times 10^{-4}$; c-hind: $p = 0.180$,<br>i-front: $p = 1.16 \times 10^{-5}$; i-mid: $p = 0.199$; i-hind: $p = 0.0463$ |
| Figure S4C (right) | Two-way ANOVA with CsCh expression and leg identity as factors | ±CsCh: $p = 0.00714$; leg identity: $p = 0.0912$; interaction between leg identity and ±CsCh: $p = 0.0414$ |
| Figure S4C (right) | Post-hoc Tukey-Kramer tests comparing CsCh+ and no-CsCh for each leg | c-front: $p = 0.880$; c-mid: $p = 1.00$; c-hind: $p = 0.480$, i-front: $p = 1.00$; i-mid: $p = 0.0418$; i-hind: $p = 1.00$ |
| Figure S4D (left) | Parametric two-way ANOVA for circular data with CsCh expression and leg identity as factors | ±CsCh: $p = 1.09 \times 10^{-6}$; leg identity: $p = 1.61 \times 10^{-8}$; interaction between leg identity and ±CsCh: $p = 1.81 \times 10^{-7}$ |
| Figure S4D (left) | Post-hoc parametric Watson-Williams multisample tests for equal means, comparing ±CsCh for each leg, with Holm-Bonferroni correction for multiple comparisons | c-front: $p = 0.00403$ *; c-mid: $p = 0.0174$; c-hind: $p = 0.0763$;<br>i-front: $p = 6.44 \times 10^{-4}$ *; i-mid: $p = 6.63 \times 10^{-4}$ *; i-hind: $p = 0.0420$<br>(where * denotes significance after multiple comparisons correction, $\alpha = 0.05$) |
| Figure S4D (middle) | Parametric two-way ANOVA for circular data with depol/hyperpol and leg identity as factors | depol/hyperpol: $p = 9.82 \times 10^{-10}$; leg identity: $p = 0.412$;<br>interaction: $p = 8.14 \times 10^{-6}$ |
| Figure S4D (middle) | Post-hoc parametric Watson-Williams multisample tests for equal means, comparing depol/hyperpol for each leg, with Holm-Bonferroni correction for multiple comparisons | c-front: $p = 2.85 \times 10^{-4}$ *; c-mid: $p = 0.0123$ *; c-hind: $p = 0.552$;<br>i-front: $p = 0.00189$ *; i-mid: $p = 0.0132$ *; i-hind: $p = 0.798$ (where * denotes significance after multiple comparisons correction, $\alpha = 0.05$) |
| Figure S4D (right) | Parametric two-way ANOVA for circular data with CsCh expression and legs as factors | ±CsCh: $p = 0.176$; leg identity: $p = 0.613$; interaction between leg identity and ±CsCh: $p = 0.232$ |
| Figure S5A | Paired t-test comparing walking and not walking | DNa02: $p = 0.00229$; DNg13: $4.59 \times 10^{-5}$ |

## References

1. Brown, T.G., and Sherrington, C.S. (1997). The intrinsic factors in the act of progression in the mammal. Proc. R. Soc. Lond. B Biol. Sci. 84, 308–319.

2. Grillner, S. (2011). Control of locomotion in bipeds, tetrapods, and fish. In Handbook of Physiology: The Nervous System: Motor Control (John Wiley & Sons, Inc.).

3. Wilson, D.M., and Wyman, R.J. (1965). Motor output patterns during random and rhythmic stimulation of locust thoracic ganglia. Biophys. J. 5, 121–143.

4. Graham, D. (1979). Effects of circum-oesophageal lesion on the behaviour of the stick insect Carausius morosus. Biol. Cybern. 32, 139–145.

5. Ryckebusch, S., and Laurent, G. (1993). Rhythmic patterns evoked in locust leg motor neurons by the muscarinic agonist pilocarpine. J. Neurophysiol. 69, 1583–1595.

6. Yellman, C., Tao, H., He, B., and Hirsh, J. (1997). Conserved and sexually dimorphic behavioral responses to biogenic amines in decapitated Drosophila. Proc. Natl. Acad. Sci. U. S. A. 94, 4131–4136.

7. Gal, R., and Libersat, F. (2006). New vistas on the initiation and maintenance of insect motor behaviors revealed by specific lesions of the head ganglia. J. Comp. Physiol. A Neuroethol. Sens. Neural Behav. Physiol. 192, 1003–1020.

8. Jordan, L.M., and Sławińska, U. (2014). Chapter 17 - The Brain and Spinal Cord Networks Controlling Locomotion. In Neuronal Networks in Brain Function, CNS Disorders, and Therapeutics, C. L. Faingold and H. Blumenfeld, eds. (Academic Press), pp. 215–233.

9. Grillner, S., and El Manira, A. (2020). Current principles of motor control, with special reference to vertebrate locomotion. Physiol. Rev. 100, 271–320.

10. Emanuel, S., Kaiser, M., Pflueger, H.-J., and Libersat, F. (2020). On the Role of the Head Ganglia in Posture and Walking in Insects. Front. Physiol. 11, 135.

11. Ijspeert, A.J., and Daley, M.A. (2023). Integration of feedforward and feedback control in the neuromechanics of vertebrate locomotion: a review of experimental, simulation and robotic studies. J. Exp. Biol. 226. 10.1242/jeb.245784.

12. Ferreira-Pinto, M.J., Ruder, L., Capelli, P., and Arber, S. (2018). Connecting circuits for supraspinal control of locomotion. Neuron 100, 361–374.

13. Beloozerova, I.N., and Sirota, M.G. (1993). The role of the motor cortex in the control of accuracy of locomotor movements in the cat. J. Physiol. 461, 1–25.

14. Drew, T., Andujar, J.-E., Lajoie, K., and Yakovenko, S. (2008). Cortical mechanisms involved in visuomotor coordination during precision walking. Brain Res. Rev. 57, 199–211.

15. Lajoie, K., and Drew, T. (2007). Lesions of area 5 of the posterior parietal cortex in the cat produce errors in the accuracy of paw placement during visually guided locomot. J. Neurophysiol. 97, 2339–2354.

16. Strauss, R., and Heisenberg, M. (1993). A higher control center of locomotor behavior in the Drosophila brain. J. Neurosci. 13, 1852–1861.

17. Drew, T., and Rossignol, S. (1984). Phase-dependent responses evoked in limb muscles by stimulation of medullary reticular formation during locomotion in thalamic cats. J. Neurophysiol. 52, 653–675.

18. Bretzner, F., and Drew, T. (2005). Contribution of the motor cortex to the structure and the timing of hindlimb locomotion in the cat: a microstimulation study. J. Neurophysiol. 94, 657–672.

19. Rho, M.J., Lavoie, S., and Drew, T. (1999). Effects of red nucleus microstimulation on the locomotor pattern and timing in the intact cat: a comparison with the motor cortex. J. Neurophysiol. 81, 2297–2315.

20. Cregg, J.M., Leiras, R., Montalant, A., Wanken, P., Wickersham, I.R., and Kiehn, O. (2020). Brainstem neurons that command mammalian locomotor asymmetries. Nat. Neurosci. 23, 730–740.

21. Josset, N., Roussel, M., Lemieux, M., Lafrance-Zoubga, D., Rastqar, A., and Bretzner, F. (2018). Distinct Contributions of Mesencephalic Locomotor Region Nuclei to Locomotor Control in the Freely Behaving Mouse. Curr. Biol. 28, 884–901.e3.

22. Bizzi, E., and Ajemian, R. (2020). From motor planning to execution: a sensorimotor loop perspective. J. Neurophysiol. 124, 1815–1823.

23. Dorkenwald, S., Matsliah, A., Sterling, A.R., Schlegel, P., Yu, S.-C., McKellar, C.E., Lin, A., Costa, M., Eichler, K., Yin, Y., et al. (2023). Neuronal wiring diagram of an adult brain. bioRxiv. 10.1101/2023.06.27.546656.

24. Scheffer, L.K., Xu, C.S., Januszewski, M., Lu, Z., Takemura, S.-Y., Hayworth, K.J., Huang, G.B., Shinomiya, K., Maitlin-Shepard, J., Berg, S., et al. (2020). A connectome and analysis of the adult Drosophila central brain. eLife 9. 10.7554/eLife.57443.

25. Phelps, J.S., Hildebrand, D.G.C., Graham, B.J., Kuan, A.T., Thomas, L.A., Nguyen, T.M., Buhmann, J., Azevedo, A.W., Sustar, A., Agrawal, S., et al. (2021). Reconstruction of motor control circuits in adult Drosophila using automated transmission electron microscopy. Cell 184, 759–774.e18.

26. Takemura, S.-Y., Hayworth, K.J., Huang, G.B., Januszewski, M., Lu, Z., Marin, E.C., Preibisch, S., Shan Xu, C., Bogovic, J., Champion, A.S., et al. (2023). A connectome of the male Drosophila ventral nerve cord. bioRxiv, 2023.06.05.543757. 10.1101/2023.06.05.543757.

27. Zheng, Z., Lauritzen, J.S., Perlman, E., Robinson, C.G., Nichols, M., Milkie, D., Torrens, O., Price, J., Fisher, C.B., Sharifi, N., et al. (2018). A complete electron microscopy volume of the brain of adult Drosophila melanogaster. Cell 174, 730–743.e22.

28. Namiki, S., Dickinson, M.H., Wong, A.M., Korff, W., and Card, G.M. (2018). The functional organization of descending sensory-motor pathways in *Drosophila*. eLife 7, e34272.

29. Bidaye, S.S., Machacek, C., Wu, Y., and Dickson, B.J. (2014). Neuronal control of *Drosophila* walking direction. Science 344, 97.

30. Chen, C.-L., Hermans, L., Viswanathan, M.C., Fortun, D., Aymanns, F., Unser, M., Cammarato, A., Dickinson, M.H., and Ramdya, P. (2018). Imaging neural activity in the ventral nerve cord of behaving adult *Drosophila*. Nat. Commun. 9, 4390.

31. Rayshubskiy, A., Holtz, S.L., D’Alessandro, I., Li, A.A., Vanderbeck, Q.X., Haber, I.S., Gibb, P.W., and Wilson, R.I. (2020). Neural circuit mechanisms for steering control in walking Drosophila. bioRxiv 2020.04.04.024703; 10.1101/2020.04.04.024703.

32. Cande, J., Namiki, S., Qiu, J., Korff, W., Card, G.M., Shaevitz, J.W., Stern, D.L., and Berman, G.J. (2018). Optogenetic dissection of descending behavioral control in Drosophila. eLife 7. 10.7554/eLife.34275.

33. Zacarias, R., Namiki, S., Card, G.M., Vasconcelos, M.L., and Moita, M.A. (2018). Speed dependent descending control of freezing behavior in *Drosophila melanogaster*. Nat. Commun. 9, 3697.

34. Bidaye, S.S., Laturney, M., Chang, A.K., Liu, Y., Bockemühl, T., Büschges, A., and Scott, K. (2020). Two brain pathways initiate distinct forward walking programs in Drosophila. Neuron 108, 469–485.e8.

35. Isaacman-Beck, J., Paik, K.C., Wienecke, C.F.R., Yang, H.H., Fisher, Y.E., Wang, I.E., Ishida, I.G., Maimon, G., Wilson, R.I., and Clandinin, T.R. (2020). SPARC enables genetic manipulation of precise proportions of cells. Nat. Neurosci. 23, 1168–1175.

36. Strauss, R., and Heisenberg, M. (1990). Coordination of legs during straight walking and turning in Drosophila melanogaster. J. Comp. Physiol. A 167, 403–412.

37. DeAngelis, B.D., Zavatone-Veth, J.A., and Clark, D.A. (2019). The manifold structure of limb coordination in walking *Drosophila*. eLife 8. 10.7554/eLife.46409.

38. Mu, L., and Ritzmann, R.E. (2005). Kinematics and motor activity during tethered walking and turning in the cockroach, Blaberus discoidalis. J. Comp. Physiol. A Neuroethol. Sens. Neural Behav. Physiol. 191, 1037–1054.

39. Jindrich, D.L., and Full, R.J. (1999). Many-legged maneuverability: dynamics of turning in hexapods. J. Exp. Biol. 202 (Pt 12), 1603–1623.

40. Graham, D. (1972). A behavioural analysis of the temporal organisation of walking movements in the 1st instar and adult stick insect (Carausius morosus). J. Comp. Physiol. 81, 23–52.

41. Franklin, R., Bell, W.J., and Jander, R. (1981). Rotational locomotion by the cockroach Blattella germanica. J. Insect Physiol. 27, 249–255.

42. Frantsevich, L.I., and Mokrushov, P.A. (1980). Turning and righting in Geotrupes (Coleoptera, Scarabaeidae). J. Comp. Physiol. 136, 279–289.

43. Zollikofer, C. (1994). Stepping patterns in ants - influence of speed and curvature. J. Exp. Biol. 192, 95–106.

44. Zolotov, V., Frantsevich, L., and Falk, E.-M. (1975). Kinematik der phototaktischen Drehung bei der Honigbiene Apis mellifera L. J. Comp. Physiol. 97, 339–353.

45. Dürr, V., and Ebeling, W. (2005). The behavioural transition from straight to curve walking: kinetics of leg movement parameters and the initiation of turning. J. Exp. Biol. 208, 2237–2252.

46. Dürr, V. (2005). Context-dependent changes in strength and efficacy of leg coordination mechanisms. J. Exp. Biol. 208, 2253–2267.

47. Frigon, A. (2017). The neural control of interlimb coordination during mammalian locomotion. J. Neurophysiol. 117, 2224–2241.

48. Sen, R., Wu, M., Branson, K., Robie, A., Rubin, G.M., and Dickson, B.J. (2017). Moonwalker descending neurons mediate visually evoked retreat in *Drosophila*. Curr. Biol. 27, 766–771.

49. Mishima, T., and Kanzaki, R. (1999). Physiological and morphological characterization of olfactory descending interneurons of the male silkworm moth, Bombyx mori. Journal of Comparative Physiology A 184, 143–160.

50. Zorović, M., and Hedwig, B. (2011). Processing of species-specific auditory patterns in the cricket brain by ascending, local, and descending neurons during standing and walking. J. Neurophysiol. 105, 2181–2194.

51. Schnell, B., Ros, I.G., and Dickinson, M.H. (2017). A descending neuron correlated with the rapid steering maneuvers of flying *Drosophila*. Curr. Biol. 27, 1200–1205.

52. Namiki, S., Ros, I.G., Morrow, C., Rowell, W.J., Card, G.M., Korff, W., and Dickinson, M.H. (2022). A population of descending neurons that regulates the flight motor of *Drosophila*. Curr. Biol. 32, 1189–1196.e6.

53. Suver, M.P., Huda, A., Iwasaki, N., Safarik, S., and Dickinson, M.H. (2016). An array of descending visual interneurons encoding self-Motion in *Drosophila*. J. Neurosci. 36, 11768–11780.

54. Fagerstedt, P., Orlovsky, G.N., Deliagina, T.G., Grillner, S., and Ullen, F. (2001). Lateral turns in the Lamprey. II. Activity of reticulospinal neurons during the generation of fictive turns. J. Neurophysiol. 86, 2257–2265.

55. Huang, K.-H., Ahrens, M.B., Dunn, T.W., and Engert, F. (2013). Spinal projection neurons control turning behaviors in zebrafish. Curr. Biol. 23, 1566–1573.

56. Aymanns, F., Chen, C.-L., and Ramdya, P. (2022). Descending neuron population dynamics during odor-evoked and spontaneous limb-dependent behaviors. eLife 11. 10.7554/eLife.81527.

57. Katsov, A.Y., Freifeld, L., Horowitz, M., Kuehn, S., and Clandinin, T.R. (2017). Dynamic structure of locomotor behavior in walking fruit flies. eLife 6. 10.7554/eLife.26410.

58. Robie, A.A., Hirokawa, J., Edwards, A.W., Umayam, L.A., Lee, A., Phillips, M.L., Card, G.M., Korff, W., Rubin, G.M., Simpson, J.H., et al. (2017). Mapping the neural substrates of behavior. Cell 170, 393–406.e28.

59. York, R.A., Brezovec, L.E., Coughlan, J., Herbst, S., Krieger, A., Lee, S.-Y., Pratt, B., Smart, A.D., Song, E., Suvorov, A., et al. (2022). The evolutionary trajectory of drosophilid walking. Curr. Biol. 32, 3005–3015.e6.

60. Kabra, M., Lee, A., Robie, A., Egnor, R., Huston, S., Rodriguez, I.F., Edwards, A., and Branson, K. APT: Animal Part Tracker v0.3.4 10.5281/zenodo.6366082.

61. Hulse, B.K., Haberkern, H., Franconville, R., Turner-Evans, D.B., Takemura, S.-Y., Wolff, T., Noorman, M., Dreher, M., Dan, C., Parekh, R., et al. (2021). A connectome of the Drosophila central complex reveals network motifs suitable for flexible navigation and context-dependent action selection. eLife 10. 10.7554/eLife.66039.

62. Westeinde, E.A., Kellogg, E., Dawson, P.M., Lu, J., Hamburg, L., Midler, B., Druckmann, S., and Wilson, R.I. (2022). Transforming a head direction signal into a goal-oriented steering command. bioRxiv, 2022.11.10.516039. 10.1101/2022.11.10.516039.

63. Mussells Pires, P., Abbott, L.F., and Maimon, G. (2022). Converting an allocentric goal into an egocentric steering signal. bioRxiv, 2022.11.10.516026. 10.1101/2022.11.10.516026.

64. Braun, J., Hurtak, F., Wang-Chen, S., and Ramdya, P. (2023). Networks of descending neurons transform command-like signals into population-based behavioral control. bioRxiv, 2023.09.11.557103. 10.1101/2023.09.11.557103.

65. Mendes, C.S., Bartos, I., Akay, T., Márka, S., and Mann, R.S. (2013). Quantification of gait parameters in freely walking wild type and sensory deprived *Drosophila melanogaster*. eLife 2, e00231.

66. Chun, C., Biswas, T., and Bhandawat, V. (2021). Drosophila uses a tripod gait across all walking speeds, and the geometry of the tripod is important for speed control. eLife 10. 10.7554/eLife.65878.

67. Chen, C.-L., Aymanns, F., Minegishi, R., Matsuda, V.D.V., Talabot, N., Günel, S., Dickson, B.J., and Ramdya, P. (2023). Ascending neurons convey behavioral state to integrative sensory and action selection brain regions. Nat. Neurosci. 26, 682–695.

68. Tsubouchi, A., Yano, T., Yokoyama, T.K., Murtin, C., Otsuna, H., and Ito, K. (2017). Topological and modality-specific representation of somatosensory information in the fly brain. Science 358, 615–623.

69. Cheong, H.S.J., Eichler, K., Stuerner, T., Asinof, S.K., Champion, A.S., Marin, E.C., Oram, T.B., Sumathipala, M., Venkatasubramanian, L., Namiki, S., et al. (2023). Transforming descending input into behavior: The organization of premotor circuits in the Drosophila Male Adult Nerve Cord connectome. bioRxiv, 2023.06.07.543976. 10.1101/2023.06.07.543976.

70. Azevedo, A., Lesser, E., Mark, B., Phelps, J., Elabbady, L., Kuroda, S., Sustar, A., Moussa, A., Kandelwal, A., Dallmann, C.J., et al. (2022). Tools for comprehensive reconstruction and analysis of Drosophila motor circuits. bioRxiv, 2022.12.15.520299. 10.1101/2022.12.15.520299.

71. Lacin, H., Chen, H.-M., Long, X., Singer, R.H., Lee, T., and Truman, J.W. (2019). Neurotransmitter identity is acquired in a lineage-restricted manner in the Drosophila CNS. elife 8. 10.7554/eLife.43701.

72. Eckstein, N., Bates, A.S., Champion, A., Du, M., Yin, Y., Schlegel, P., Lu, A.K., Rymer, T., Finley-May, S., Paterson, T., et al. (2023). Neurotransmitter classification from electron microscopy images at synaptic sites in Drosophila melanogaster. bioRxiv, https://www.biorxiv.org/content/10.1101/2020.06.12.148775v3.

73. Lu, J., Behbehani, A., Hamburg, L., Westeinde, E.A., Dawson, P.M., Lyu, C., Maimon, G., Dickinson, M., Druckmann, S., and Wilson, R.I. (2021). Transforming representations of movement from body- to world-centric space. Nature 601, 98–104.

74. Lyu, C., Abbott, L.F., and Maimon, G. (2021). Building an allocentric travelling direction signal via vector computation. Nature 601, 92–97.

75. Stone, T., Webb, B., Adden, A., Weddig, N.B., Honkanen, A., Templin, R., Wcislo, W., Scimeca, L., Warrant, E., and Heinze, S. (2017). An anatomically constrained model for path integration in the bee brain. Curr. Biol. 27, 3069–3085.e11.

76. Seelig, J.D., and Jayaraman, V. (2015). Neural dynamics for landmark orientation and angular path integration. Nature 521, 186–191.

77. Green, J., Vijayan, V., Mussells Pires, P., Adachi, A., and Maimon, G. (2019). A neural heading estimate is compared with an internal goal to guide oriented navigation. Nat. Neurosci. 22, 1460–1468.

78. Geurten, B.R.H., Jähde, P., Corthals, K., and Göpfert, M.C. (2014). Saccadic body turns in walking Drosophila. Front. Behav. Neurosci. 8, 365.

79. Cruz, T.L., Pérez, S.M., and Chiappe, M.E. (2021). Fast tuning of posture control by visual feedback underlies gaze stabilization in walking Drosophila. Current Biology. 10.1016/j.cub.2021.08.041.

80. Drew, T., Dubuc, R., and Rossignol, S. (1986). Discharge patterns of reticulospinal and other reticular neurons in chronic, unrestrained cats walking on a treadmill. J. Neurophysiol. 55, 375–401.

81. Garcia-Rill, E., Skinner, R.D., and Fitzgerald, J.A. (1983). Activity in the mesencephalic locomotor region during locomotion. Exp. Neurol. 82, 609–622.

82. Orlovsky, G.N. (1970). Work of the reticulo-spinal neurones during locomotion. Biophysics.

83. Kasicki, S., Grillner, S., Ohta, Y., Dubuc, R., and Brodin, L. (1989). Phasic modulation of reticulospinal neurones during fictive locomotion and other types of spinal motor activity in lamprey. Brain Res. 484, 203–216.

84. Armstrong, D.M., and Drew, T. (1984). Discharges of pyramidal tract and other motor cortical neurones during locomotion in the cat. J. Physiol. 346, 471–495.

85. Armstrong, D.M., and Edgley, S.A. (1984). Discharges of Purkinje cells in the paravermal part of the cerebellar anterior lobe during locomotion in the cat. J. Physiol. 352, 403–424.

86. Matsuyama, K., and Drew, T. (2000). Vestibulospinal and reticulospinal neuronal activity during locomotion in the intact cat. I. Walking on a level surface. J. Neurophysiol. 84, 2237–2256.

87. Deliagina, T.G., Zelenin, P.V., Fagerstedt, P., Grillner, S., and Orlovsky, G.N. (2000). Activity of reticulospinal neurons during locomotion in the freely behaving lamprey. J. Neurophysiol. 83, 853–863.

88. Oueghlani, Z., Simonnet, C., Cardoit, L., Courtand, G., Cazalets, J.-R., Morin, D., Juvin, L., and Barrière, G. (2018). Brainstem steering of locomotor activity in the newborn rat. J. Neurosci. 38, 7725–7740.

89. Kozlov, A.K., Kardamakis, A.A., Hellgren Kotaleski, J., and Grillner, S. (2014). Gating of steering signals through phasic modulation of reticulospinal neurons during locomotion. Proc. Natl. Acad. Sci. U. S. A. 111, 3591–3596.

90. Fujiwara, T., Brotas, M., and Chiappe, M.E. (2022). Walking strides direct rapid and flexible recruitment of visual circuits for course control in Drosophila. Neuron 110, 2124–2138.e8.

91. Wosnitza, A., Bockemühl, T., Dübbert, M., Scholz, H., and Büschges, A. (2013). Inter-leg coordination in the control of walking speed in Drosophila. J. Exp. Biol. 216, 480–491.

92. Drew, T. (1991). Functional organization within the medullary reticular formation of the intact unanesthetized cat. III. Microstimulation during locomotion. J. Neurophysiol. 66, 919–938.

93. Bidaye, S.S., Bockemühl, T., and Büschges, A. (2018). Six-legged walking in insects: how CPGs, peripheral feedback, and descending signals generate coordinated and adaptive motor rhythms. J. Neurophysiol. 119, 459–475.

94. Azevedo, A.W., Dickinson, E.S., Gurung, P., Venkatasubramanian, L., Mann, R.S., and Tuthill, J.C. (2020). A size principle for recruitment of *Drosophila* leg motor neurons. eLife 9. 10.7554/eLife.56754.

95. Dickinson, M.H., Farley, C.T., Full, R.J., Koehl, M.A., Kram, R., and Lehman, S. (2000). How animals move: an integrative view. Science 288, 100–106.

96. Full, R.J., Blickhan, R., and Ting, L.H. (1991). Leg design in hexapedal runners. J. Exp. Biol. 158, 369–390.

97. Heinze, S., Narendra, A., and Cheung, A. (2018). Principles of insect path integration. Curr. Biol. 28, R1043–R1058.

98. Grob, R., el Jundi, B., and Fleischmann, P.N. (2021). Towards a common terminology for arthropod spatial orientation. Ethol. Ecol. Evol. 33, 338–358.

99. Hulse, B.K., and Jayaraman, V. (2020). Mechanisms underlying the neural computation of head direction. Annu. Rev. Neurosci. 43, 31–54.

100. Wilson, R.I. (2023). Neural networks for navigation: From connections to computations. Annu. Rev. Neurosci. 46, 403–423.

101. Vyas, S., Golub, M.D., Sussillo, D., and Shenoy, K.V. (2020). Computation through neural population dynamics. Annu. Rev. Neurosci. 43, 249–275.

102. Gallego, J.A., Perich, M.G., Miller, L.E., and Solla, S.A. (2017). Neural manifolds for the control of movement. Neuron 94, 978–984.

103. Shepherd, D., Sahota, V., Court, R., Williams, D.W., and Truman, J.W. (2019). Developmental organization of central neurons in the adult Drosophila ventral nervous system. J. Comp. Neurol. 527, 2573–2598.

104. Harris, R.M., Pfeiffer, B.D., Rubin, G.M., and Truman, J.W. (2015). Neuron hemilineages provide the functional ground plan for the Drosophila ventral nervous system. elife 4. 10.7554/eLife.04493.

105. Silies, M., Gohl, D.M., Fisher, Y.E., Freifeld, L., Clark, D.A., and Clandinin, T.R. (2013). Modular use of peripheral input channels tunes motion-detecting circuitry. Neuron 79, 111–127.

106. Pfeiffer, B., Ngo, T.-T.B., Hibbard, K.L., Murphy, C., Jenett, A., Truman, J.W., and Rubin, G.M. (2010). Refinement of tools for targeted gene expression in *Drosophila*. Genetics 186, 735–755.

107. Pearn, M.T., Randall, L.L., Shortridge, R.D., Burg, M.G., and Pak, W.L. (1996). Molecular, biochemical, and electrophysiological characterization of Drosophila norpA mutants. J. Biol. Chem. 271, 4937–4945.

108. Moore, R.J., Taylor, G.J., Paulk, A.C., Pearson, T., van Swinderen, B., and Srinivasan, M.V. (2014). FicTrac: a visual method for tracking spherical motion and generating fictive animal paths. J. Neurosci. Methods 225, 106–119.

109. Pnevmatikakis, E.A., and Giovannucci, A. (2017). NoRMCorre: An online algorithm for piecewise rigid motion correction of calcium imaging data. J. Neurosci. Methods 291, 83–94.

110. Bates, A.S., Manton, J.D., Jagannathan, S.R., Costa, M., Schlegel, P., Rohlfing, T., and Jefferis, G.S. (2020). The natverse, a versatile toolbox for combining and analysing neuroanatomical data. eLife 9. 10.7554/elife.53350.

111. Berens, P. (2009). CircStat: A MATLAB Toolbox for Circular Statistics. J. Stat. Softw. 31, 1–21.

112. Drosophila Information Service, Volume 60 (1984) https://www.ou.edu/journals/dis/DIS60/DIS60.html.

113. Pologruto, T.A., Sabatini, B.L., and Svoboda, K. (2003). ScanImage: flexible software for operating laser scanning microscopes. Biomed. Eng. Online 2, 13.

114. Gouwens, N.W., and Wilson, R.I. (2009). Signal propagation in Drosophila central neurons. J. Neurosci. 29, 6239–6249.

115. Buhmann, J., Sheridan, A., Malin-Mayor, C., Schlegel, P., Gerhard, S., Kazimiers, T., Krause, R., Nguyen, T.M., Heinrich, L., Lee, W.-C.A., et al. (2021). Automatic detection of synaptic partners in a whole-brain *Drosophila* electron microscopy data set. Nature Methods 18, 771–774. 10.1038/s41592-021-01183-7.

116. Heinrich, L., Funke, J., Pape, C., Nunez-Iglesias, J., and Saalfeld, S. (2018). Synaptic Cleft Segmentation in Non-isotropic Volume Electron Microscopy of the Complete Drosophila Brain. In Medical Image Computing and Computer Assisted Intervention – MICCAI 2018 (Springer International Publishing), pp. 317–325.

117. Ito, K., Shinomiya, K., Ito, M., Armstrong, J.D., Boyan, G., Hartenstein, V., Harzsch, S., Heisenberg, M., Homberg, U., Jenett, A., et al. (2014). A systematic nomenclature for the insect brain. Neuron 81, 755–765.

118. Lesser, E., Azevedo, A.W., Phelps, J.S., Elabbady, L., Cook, A., Mark, B., Kuroda, S., Sustar, A., Moussa, A., Dallmann, C.J., et al. (2023). Synaptic architecture of leg and wing motor control networks in Drosophila. bioRxiv. 10.1101/2023.05.30.542725.

119. French, A.S. (1976). Practical nonlinear system analysis by Wiener kernel estimation in the frequency domain. Biol. Cybern. 24, 111–119.

120. Liu, W.W., and Wilson, R.I. (2013). Glutamate is an inhibitory neurotransmitter in the Drosophila olfactory system. Proc. Natl. Acad. Sci. U. S. A. 110, 10294–10299.

121. Cully, D.F., Paress, P.S., Liu, K.K., Schaeffer, J.M., and Arena, J.P. (1996). Identification of a Drosophila melanogaster glutamate-gated chloride channel sensitive to the antiparasitic agent avermectin. J. Biol. Chem. 271, 20187–20191.

122. Allen, A.M., Neville, M.C., Birtles, S., Croset, V., Treiber, C.D., Waddell, S., and Goodwin, S.F. (2020). A single-cell transcriptomic atlas of the adult Drosophila ventral nerve cord. eLife 9. 10.7554/eLife.54074.

